# Quartz-Seq2: a high-throughput single-cell RNA-sequencing method that effectively uses limited sequence reads

**DOI:** 10.1101/159384

**Authors:** Yohei Sasagawa, Hiroki Danno, Hitomi Takada, Masashi Ebisawa, Kaori Tanaka, Tetsutaro Hayashi, Akira Kurisaki, Itoshi Nikaido

## Abstract

High-throughput single-cell RNA-seq methods assign limited unique molecular identifier (UMI) counts as gene expression values to single cells from shallow sequence reads and detect limited gene counts. We thus developed a high-throughput single-cell RNA-seq method, Quartz-Seq2, to overcome these issues. Our improvements in the reaction steps make it possible to effectively convert initial reads to UMI counts (at a rate of 30%–50%) and detect more genes. To demonstrate the power of Quartz-Seq2, we analyzed approximately 10,000 transcriptomes in total from *in vitro* embryonic stem cells and an *in vivo* stromal vascular fraction with a limited number of reads.

## Background

Single-cell transcriptome analysis is a powerful tool to identify nongenetic cellular heterogeneity, which includes differences in cell type due to differentiation and differences in cell state within a cell population. In previous studies, various methods for single-cell RNA-seq were developed [1
–19]. Some of these that generate read coverage across all transcripts have been exploited to detect alternative transcription splicing isoforms [14,17], and others using a unique molecular identifier (UMI) have been applied to quantify the number of transcripts expressed in a cell [1–3,6,7,9,11–13,20]. To extract substantial information on a cell population, such as the composition of different cell types or the distribution of cell states, it is necessary to analyze hundreds or thousands of cells. Cell barcoding is a key technology for this, which enables us to deal with samples from numerous cells in a single tube. Cell barcoding technology, which tags nucleotides unique to each cell to target RNA molecules from that cell, is a key technology for increasing the throughput of single-cell RNA-seq [16,18]. Mixing cDNA tagged with cell barcodes before whole-transcript amplification decreases the cost of reaction reagents and the laboriousness of experimental steps. There are two types of cell barcoding technology according to the method of cell sampling used. One method involves single cells being selectively sorted to multi-well plates using flow cytometry, which allows us to remove dead or aggregated cells. Besides, transcriptome data can be linked to cellular information obtained by flow cytometry. The other method involves single cells and barcoded beads being captured in water-in-oil droplets using droplet-generation microfluidic devices [6,7]. In this latter method, thousands of cells can probabilistically be captured in half an hour. However, the total number of sequence reads generated by a deep sequencer is still limited. To increase the number of analyzed cells, each cell is assigned a limited number of initial sequence reads. For example, approximately 400 million initial fastq reads were sequenced for 3,000–4,000 cells in several previous studies [6,7]. In this case, the input data size for a single cell involves shallow initial reads (100,000 fastq reads) (Figure 1b). UMI counts were converted from shallow initial sequence reads for each cell, and the conversion ratio of that was limited to approximately one-tenth [21]. Ideally, greater UMI counts should be generated from limited sequence reads because the increase in UMI count assigned to each cell leads to the detection of low-copy genes and the identification of cell-type-specific genes using statistical tests.

**Figure 1.**
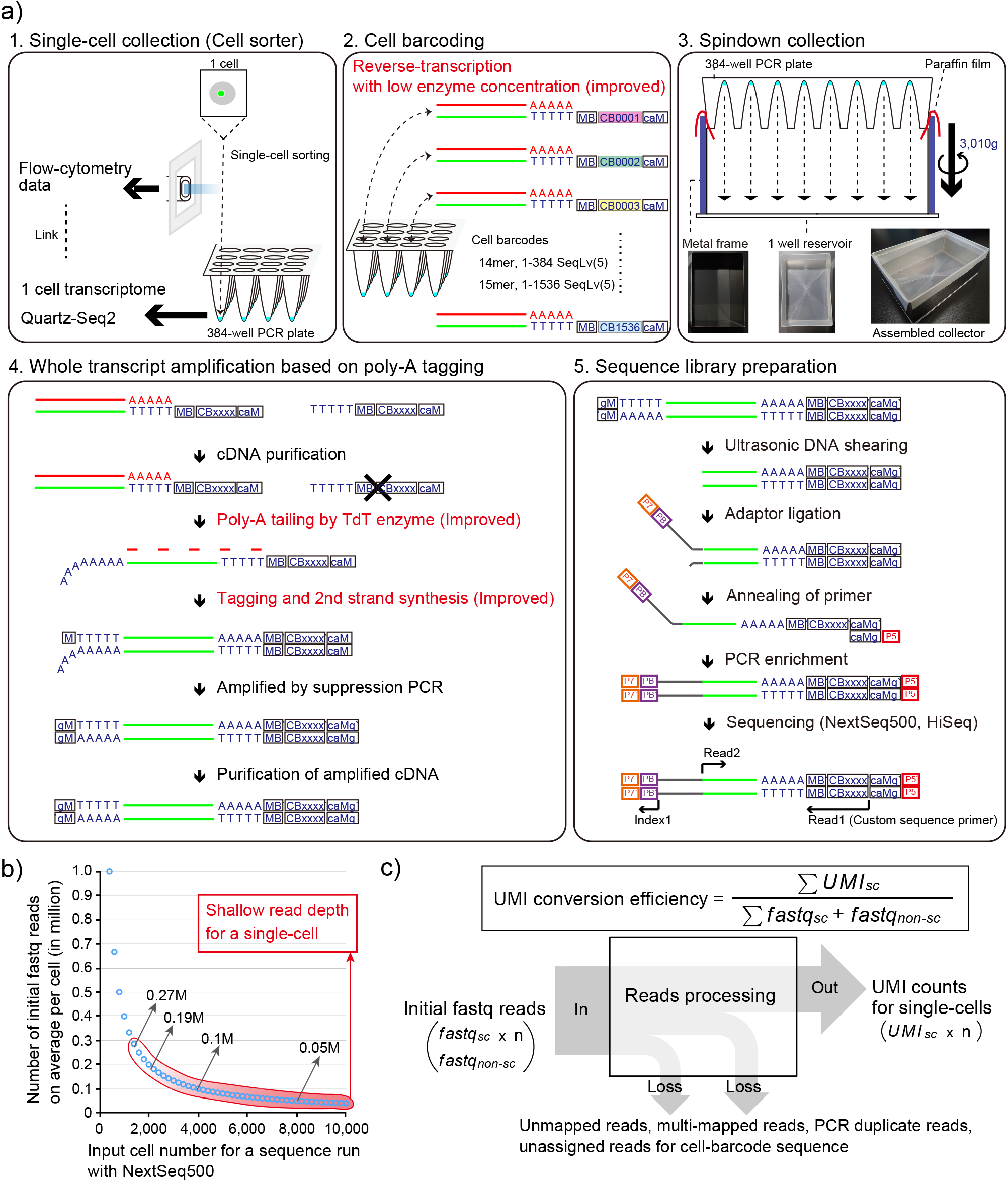
Overview of Quartz-Seq2 experimental processes. a) Quartz-Seq2 consisted of five steps. (1) Each single cell in a droplet was sorted into lysis buffer in each well of a 384-well PCR plate using flow cytometry analysis data. (2) Poly-adenylated RNA in each well was reverse-transcribed into first-strand cDNA with reverse transcription primer, which had a unique cell barcode. We prepared 384 or 1,536 kinds of cell barcode (CB) with a unique sequence based on the Sequence–Levenshtein distance (SeqLv). The edit distance of SeqLv was 5. RT primer also had a UMI sequence for reduction of PCR bias (MB) and a poly-dT sequence for binding to poly-A RNA. (3) Cell-barcode-labeled cDNAs from all 384 wells were promptly collected by centrifugation using assembled collectors. (4) Collected first-strand cDNAs were purified and concentrated for subsequent whole-transcript amplification. In the poly-A tailing step, purified cDNA was extended with a poly-A tail by terminal deoxynucleotidyl transferase (TdT). Subsequently, second-strand cDNA was synthesized with a tagging primer, which had a poly-dT sequence. The resulting second-strand cDNA had a PCR primer sequence (M) at both ends of it. The cDNA was amplifiable in subsequent PCR amplification. (5) For conversion from amplified cDNA to sequence library DNA, we fragmented the amplified cDNA using the ultrasonicator Covaris. Such fragmented cDNA was ligated with a truncated Y-shaped sequence adaptor, which had an Illumina flow-cell binding sequence (P7) and a pool barcode sequence (PB). PB made it possible to mix different sets of cell-barcode-labeled cDNA. Ligated cDNA, which had CB and MB sequences, was enriched by PCR amplification. The resulting sequence library DNA contained P7 and P5 flow-cell binding sequences at respective ends of the DNA. We sequenced the cell-barcode site and the UMI site at Read1, the pool-barcode site at Index1, and the transcript sequence at Read2. b) The relationship between initial fastq reads and the number of single cells for sequence analysis in NextSeq500 runs. Typically, one sequence run with NextSeq 500/550 High Output v2 Kit reads out 400–450 M fastq reads. The x-axis represents the input cell number for one sequence run. The y-axis represents the initial data size (fastq reads) on average per cell. Red circle represents the typical range of shallow input read depth for a single cell. c) We define the formula for calculating the UMI conversion efficiency. Each parameter is defined as follows: *UMI_sc_*: the number of UMI count, assigned to a single-cell sample, *fastq_sc_*: the number of fastq reads derived from each single-cell sample, *fastq_non-sc_*: the number of fastq reads derived from non-single-cell samples, which include experimental byproducts such as WTA adaptors, WTA byproducts, and non-STAMPs. Initial fastq reads are composed of *fastq_sc_* and *fastq_non-sc_*.

In this study, we developed a novel high-throughput single-cell RNA-seq method, Quartz-Seq2. As Quartz-Seq is a sensitive and reproducible single-cell RNA-seq method, Quartz-Seq2 was developed based on it [15]. Quartz-Seq is based on a poly-A tagging strategy. By the combination of molecular biological improvements including major improvement of poly-A tagging, Quartz-Seq2 resulted in an increase in the effectiveness with which the initial sequence reads were converted to the expression UMI counts (UMI conversion efficiency: 30%–50%). To demonstrate the highly effective use of initial reads in Quartz-Seq2, we analyzed a population of approximately 9,000 mouse embryonic stem (ES) cells as *in vitro* cells and approximately 1,000 cells from the stromal vascular fraction (SVF) as *in vivo* cells.

## Results

### Outline of Quartz-Seq2 experiment

To increase the UMI conversion efficiency (Figure 1), we improved several steps in the preparation of the single-cell RNA-seq library (Additional file 1: Figure S1), resulting in the development of Quartz-Seq2. Below, we explain the five steps of the Quartz-Seq2 procedure (Figure 1).

1. The first step is single-cell collection using a cell sorter. We selectively sort living single cells into lysis buffer in a 384-well PCR plate without dead cells. In cell sorting, various types of channel information, such as the intensity of fluorescence, are obtained for each cell. This enables us to link the transcriptome to cellular information from a cell sorter for each cell.
2. The second step is cell barcoding. Each well contains lysis buffer and reverse-transcription (RT) primer, which includes a cell barcode sequence (14- or 15-mer), a UMI sequence (8-mer), and an oligo-dT sequence (24-mer). Using these RT primers, respective RNA from single cells is converted to cDNA with unique cell barcodes. Note that a long RT primer resulted in a severe problem regarding the synthesis of byproducts at the downstream reaction in our system (Additional file 1: Figure S2 and Supplemental Note). Therefore, we use a relatively short RT primer (73- or 74-mer), which allows us to skip the step of removing byproducts using exonuclease I. We design two types of RT primer set (v3.1: 384 barcodes; v3.2: 1536 barcodes). Within each primer set, barcode sequences are designed such that the minimum Sequence–Levenshtein distance between two sequences should be greater than 5, which leads to the correction of mutations of two nucleotides, including substitution, insertion, or deletion in sequence reads [22]. We also optimize the buffer and temperature in the RT reaction, leading to an improvement of RT efficiency from that of the original Quartz-Seq (Additional file 1: Figure S3 and Supplemental Note). We apply a low enzyme concentration in reverse transcription to Quartz-Seq2. These conditions reduce the cost and technical variability of Quartz-Seq2. We describe the details of this in the subsection entitled “Reduction of enzyme concentration in reverse transcription decreased the experimental cost of Quartz-Seq2”.
3. The third step involves the pooling of cell-barcoded cDNA. By cell barcoding, Quartz-Seq2 can pool cDNA of up to 1,536 individual cells into one mixture. We developed a rapid and high-throughput method for collecting small volumes of cDNA in multiwell plates. This method also achieved higher efficiency of collection than dispensing with pipettes and tips (Additional file 1: Figure S4 and Supplemental Note). As the efficiency of cDNA purification after pooling was 93.77%, we estimated that approximately 80% of cell-barcoded cDNA could be used for subsequent whole-transcript amplification in our system (Additional file 1: Figure S3a and Supplemental Note).
4. The fourth step is whole-transcript amplification based on an improved poly-A tagging strategy. Poly-A tagging is one of the methods of converting first-strand cDNA to amplifiable cDNA. First-strand cDNA is extended with a poly-A tail by the terminal transferase. Subsequently, second-strand cDNA is synthesized with tagging primers that contain a poly-dT sequence, followed by PCR amplification. Here, we improve the efficiency of poly-A tagging by 3.6-fold. This improvement is a crucial point in the development of Quartz-Seq2. We describe the details of this in the next subsection.
5. The fifth step is library preparation for deep sequencing. Amplified cDNA is fragmented, ligated with the sequence adapter, and amplified by PCR. In sequencing using Illumina sequencers, a sequence for a cell barcode and a UMI is read in Read1, while a sequence for a region of a transcript (mRNA) is read in Read2.

### Improvement of poly-A tagging efficiency

We previously reported Quartz-Seq based on the poly-A tagging strategy, which has significant potential for detecting a large number of genes expressed in a cell [15]. However, the efficiency of poly-A tagging itself for single-cell RNA-seq has not been improved. We hypothesized that the improvement of poly-A tagging would lead to high UMI conversion efficiency. Therefore, we attempted to improve the efficiency of this tagging step. Poly-A tagging is composed of two processes: (1) The first-strand cDNA is modified with a poly-A tail by terminal deoxynucleotidyl transferase. (2) Next, the poly-A-tailed cDNA is annealed with a tagging primer, which has a poly-dT sequence. Then, the second-strand cDNA is extended. The resulting second-strand cDNA is amplifiable cDNA, which has a PCR primer sequence at both ends of it.

It is known that the DNA yield of amplified cDNA generally reflects the quantitative performance of single-cell RNA-seq methods [3,13,14]. Thus, we determined the effects of various buffers for the poly-A tailing step on the amplified cDNA yield. We performed the poly-A tailing reaction with various buffers using purified first-strand cDNA from 1 ng of total RNA (Figure 2a, Additional file 1: Figure S5). Finally, we obtained amplified cDNA. We found that the use of T55 buffer in the poly-A tailing reaction efficiently improved the cDNA yield (Figure 2a). The amount of amplified cDNA increased 2.88-fold using T55 buffer compared with the level using Quartz-Seq buffer (Figure 2a). In these buffer conditions, we did not observe any obvious byproducts derived from the RT primer (Figure 2a, Additional file 1: Figures S2 and S5, and Supplemental Note).

**Figure 2.**
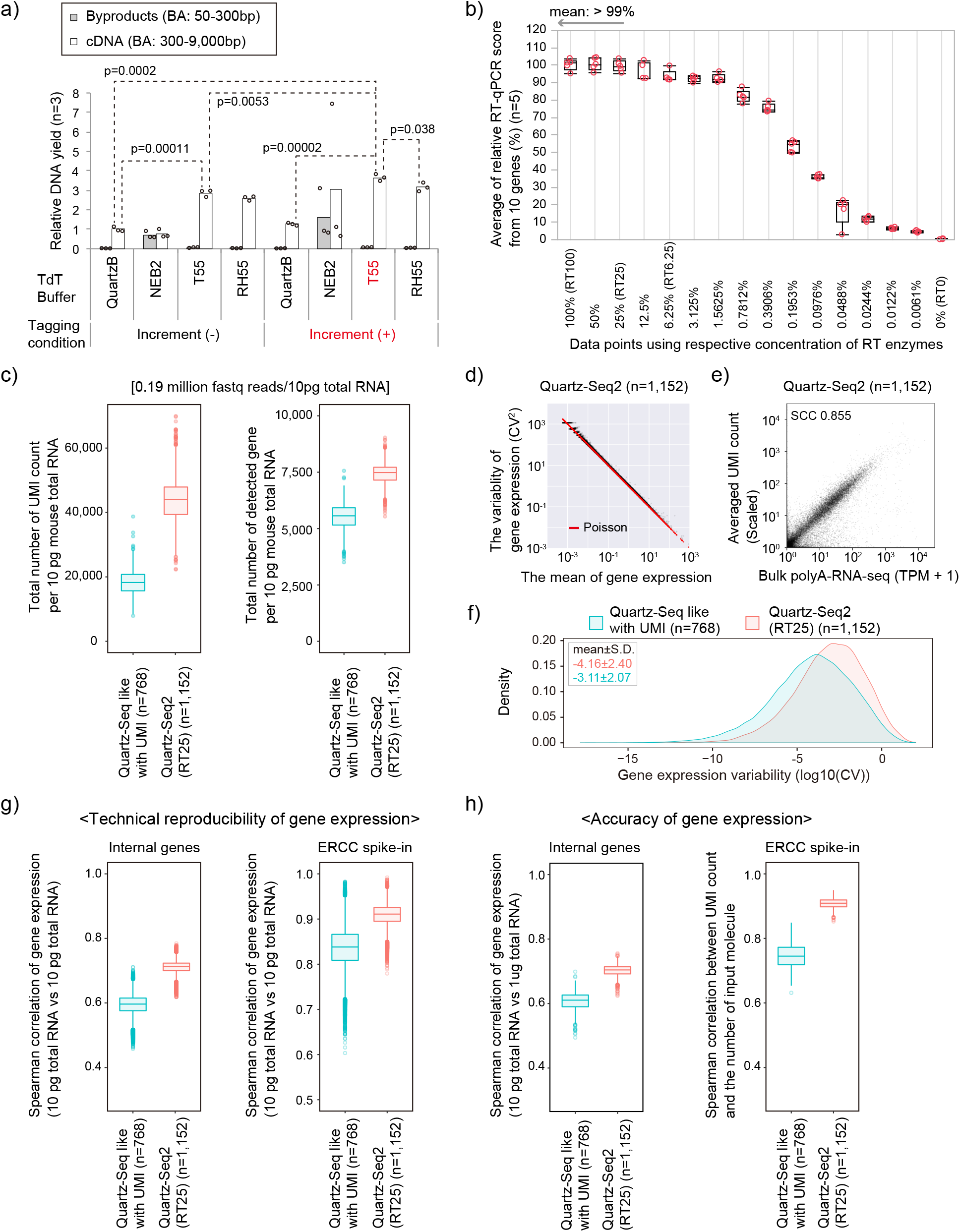
Sequence performance of Quartz-Seq2 with molecular biological improvements. a) Improvement of poly-A tagging efficiency. Bar plot represents the relative DNA yield in various poly-A tagging conditions using purified first-strand cDNA from 1 ng of total RNA. T55 buffer as the terminal deoxynucleotidyl transferase (TdT) buffer and the temperature condition “Increment” for the poly-A tagging step improved the cDNA yield of whole-transcript amplification. Buffer compositions are indicated in Additional file 3: Table S2. QuartzB represents use of a Quartz-Seq-like buffer as a positive control, in accordance with the approach described in the original Quartz-Seq paper. Finally, we quantified cDNA yield (300–9,000 bp) and byproduct DNA yield (50–300 bp) using Bioanalyzer (Agilent). The presented p-value was obtained using two-tailed Welch’s t-test. b) Reverse transcription efficiency with serially diluted RT enzymes. The x-axis represents the average relative RT qPCR score from 10 genes. Detailed concentrations of RT enzymes are presented in Additional file 1: Figure S7. c,f–h) Comparison between Quartz-Seq2 in the RT25 condition and Quartz-Seq-like conditions regarding sequence performance. c) We analyzed 384 wells with 10 pg of total RNA and used approximately 0.19 M fastq reads on average per well. We show the UMI count and gene count in box plots. d) The panel shows a scatter plot between the mean of gene expression and the variability of gene expression with 10 pg of total RNA in 384 wells. Red lines represent the theoretical variability of gene expression in the form of a Poisson distribution. e) Gene expression reproducibility between bulk poly-A-RNA-seq (1 μg of total RNA) and Quartz-Seq2 (10 pg of total RNA, averaged over 384 wells). f) Dispersion of gene expression. The x-axis represents gene expression variability. g) Reproducibility of gene expression for internal gene and external control RNA. h) Accuracy of gene expression for internal gene and external control RNA.

Next, we added an “Increment” temperature condition for the tagging and second-strand synthesis steps (see Methods). In this condition, the reaction temperature of these steps was gradually increased. As a result, the amount of cDNA tended to increase, by approximately 1.2-fold (Figure 2a). Moreover, upon combining T55 buffer and the Increment condition, the amount of cDNA increased approximately 3.6-fold. We also confirmed the reproducibility of this phenomenon of cDNA increment in additional experiments (Additional file 1: Figure S5). Moreover, we confirmed the amplified cDNA yield of various genes by qPCR analysis as another assay. Specifically, we determined the qPCR scores of eight genes from amplified cDNA and nonamplified cDNA (Additional file 1: Figure S5c). Spearman’s rank correlation coefficients (SCCs) between amplification and nonamplification were approximately 0.79 in the condition (T55+Increment). The SCC was approximately 0.66 in Quartz-Seq-like conditions. We also observed clear increments of qPCR scores for almost all genes. These results show that the combination of T55 buffer and this temperature condition improved the efficiency of the poly-A tagging step. We also found that other conditions (NBF40+Increment) improved the cDNA yield. However, under these conditions, byproducts were clearly synthesized (Additional file 1: Figures S2c and S5b). Moreover, the amount of cDNA with T55 buffer was slightly greater than that with RH55 (Figure 2 and Additional file 1: Figure S5a). Therefore, we used the combination of T55 buffer and the “Increment” temperature condition for the poly-A tagging strategy for Quartz-Seq2.

### Reduction of enzyme concentration in reverse transcription decreased the experimental cost of Quartz-Seq2

The cost of experiments for the single-cell RNA-seq method is one of the most important benchmarks regarding high-throughput performance. The cost of experimental preparation per cell was approximately ¥2,600 ($23) for our previously reported Quartz-Seq, which does not use cell barcoding (Additional file 1: Figure S6a). To improve on this value, we first applied the “RT100” enzyme condition in reverse transcription to Quartz-Seq2. In this condition, we used approximately 20U reverse transcriptase in 2 μL of solution for reverse transcription. This enzyme concentration in the RT reaction is broadly used for various molecular biological applications including single-cell RNA-seq methods [3,14,15]. By using the cell barcoding strategy, the cost of experimental preparation for Quartz-Seq2 under the “RT100” condition was reduced (¥122 or $1.08 per cell) (Additional file 1: Figure S6a).

We found that 65% of the cost of experimental steps is derived from reverse transcription in Quartz-Seq2 under the “RT100” condition (Additional file 1: Figure S7a). To further reduce the cost of experimental preparation of sequence libraries on a large scale, we investigated the effect of a low enzyme concentration in reverse transcription in Quartz-Seq2. In the assessment assay, we noticed that a low enzyme concentration did not markedly affect the efficiency of reverse transcription in T100 buffer (Additional file 1: Figure S7b). We performed a similar experiment with a broader range of concentrations of enzymes in reverse transcription under conditions with T100 buffer (Figure 2b and Additional file 1: Figure S7c). We found that the RT25 condition maintained the efficiency of reverse transcription at the 99% level on average, which was comparable to that in the RT100 condition. Therefore, we prepared three technical replicates of a 384-well PCR plate with 10 pg of total RNA with the “RT100” condition or the “RT25” low-enzyme condition. In the “RT25” condition, the cDNA yield showed a tendency for a slight increase of approximately 1.17-fold, which although not being a major improvement did at least not involve a decrease (Additional file 1: Figure 7d). The cost of experimental preparation per cell became approximately ¥46–¥63 ($0.40–$0.56) in the “RT25” condition (Figure S6a). We thus mainly used the “RT25” condition for Quartz-Seq2.

### Evaluation of the quantitative performance of Quartz-Seq2 using 10 pg of purified total RNA

Finally, we adopted three molecular biological improvements [poly-A tailing buffers (T55), the “Increment” temperature condition, and low-enzyme concentration (RT25) in reverse transcription] for Quartz-Seq2 (Additional file 1: Figure S1). To determine the technical variability and specificity of Quartz-Seq2, we performed whole-transcript amplification using 10 pg of diluted mouse total RNA as a single-cell-like averaged sample with the v3.1 384 cell-barcode RT primer in a 384-well plate (Figure 2). In this experiment, we used the “RT25” enzyme concentration for Quartz-Seq2. The effect of RT enzyme concentration on the quantitative performance is specifically described in the last paragraph in this subsection of the paper. We analyzed 10 pg of total RNA in all wells at approximately 0.19 M fastq reads on average per well. In the case of the Quartz-Seq-like conditions, we detected 18,407±4,040 UMI counts and 5,728±604 gene counts (n=768 wells from two 384-well plates). In the case of Quartz-Seq2, we achieved a UMI count of 44,100±7,521 and a gene count of 7,442±484 (n=1,152 wells from three 384-well plates)(Figure 2c). We observed similar results at the level of individual 384-well plates (Additional file 1: Figure S8a). We also calculated the UMI conversion efficiency for the respective protocol at approximately 0.19 million initial fastq reads on average per well. We defined the formula for calculating the “UMI conversion efficiency,” which indicates how effectively initial fastq reads can be converted to UMI counts (Figure 1c and Additional file 1: Figure S9). The UMI conversion efficiency levels of Quartz-Seq2 and Quartz-Seq-like were about 22.88% and 9.55%, respectively. The UMI conversion efficiency was reproducible among individual 384-well plates (Additional file 1: Figure S8a). In addition, the UMI conversion efficiency depended on the Read2 length and the number of initial fastq reads (Additional file 1: Figure S8b). These results indicated that the combination of the molecular biological improvements for Quartz-Seq2 clearly improved the conversion ratio from target RNA to sequence library DNA.

We also validated the technical reproducibility of gene expression. In the mean-CV (coefficient of variation) plot, technical gene expression variability of Quartz-Seq2 became fairly close to the theoretical variability of a Poisson distribution (Figure 2d). The mean of gene expression variability for Quartz-Seq2 was lower than that for the Quartz-Seq-like method (Figure 2f). To quantify the technical reproducibility of internal gene expression and external gene (ERCC spike-in RNA) expression, we used a pairwise comparison of technical replicates for each protocol. Regarding the reproducibility of internal gene expression, the SCCs of Quartz-Seq2 and Quartz-Seq-like were 0.71±0.01 and 0.59±0.02, respectively. Regarding the reproducibility of external gene expression, the SCCs of Quartz-Seq2 and Quartz-Seq-like were 0.90±0.02 and 0.83±0.04, respectively. Subsequently, we validated the accuracy of internal/external gene expression. We observed that the average internal gene expression of Quartz-Seq2 highly correlated with the internal gene expression of conventional RNA-seq (Figure 2e). We calculated all combinations of pairwise correlation between the internal gene expression of Quartz-Seq2 with 10 pg of total RNA and that of conventional RNA-seq with 1 μg of total RNA. Moreover, we calculated the pairwise correlation between the external gene expression of Quartz-Seq2 and the input molecule count of external genes (ERCC spike-in RNA). Regarding the accuracy of internal gene expression, the SCCs of Quartz-Seq2 and Quartz-Seq-like were 0.70±0.01 and 0.60±0.02, respectively. Regarding the accuracy of external gene expression, the SCCs of Quartz-Seq2 and Quartz-Seq-like were 0.90±0.01 and 0.74±0.03, respectively. These results indicate that the combination of the molecular biological improvements for Quartz-Seq2 clearly improved the technical reproducibility and accuracy of gene expression.

We noted that a low enzyme concentration improved the quantitative performance of Quartz-Seq2. We validated the quantitative performance with 10 pg of total RNA in the “RT100” and “RT25” conditions at various input data sizes. Unexpectedly, we found that the “RT25” condition improved the quantitative performance (Additional file 1: Figure S8a). We compared RT25 with RT100 at approximately 0.096 M fastq reads on average per well. In the case of Quartz-Seq2 in the “RT25” condition, we achieved a UMI count of 30,117±789 and a gene count of 6,320±35 (three 384-well plates). In the case of Quartz-Seq2 in the “RT100” condition, we achieved a UMI count of 25,671±1020 and a gene count of 5,889±35 (three 384-well plates). We also observed that the well-to-well technical variability for the UMI count and gene count clearly decreased in the “RT25” condition (Additional file 1: Figure S8a). These results showed that the “RT25” low-enzyme condition clearly reduced the experimental cost and improved the quantitative performance. We thus applied the “RT25” condition in subsequent experiments using real single cells. Note that, in the RT1.5625-RT6.25 condition, reverse-transcription efficiency was maintained at an average level of over 90% (Figure 2b). The average reverse-transcription efficiency rapidly decreased below the RT1.56 low enzyme concentration. In addition, in actual experiments (not serially diluted experiments) at conditions below RT3.12, it was not guaranteed that enzymes could be collected given the viscosity and low volume of the mixture. Therefore, we validated the RT6.25 condition for subsequent analysis. It seems that UMI counts, gene counts, and ERCC capture efficiency (also called as “ERCC spike-in RNA detection efficiency”) slightly increased in the RT6.25 condition, which although not being a major improvement did at least not involve a decrease (Additional file 1: Figure S10). The cost of experimental preparation per cell became approximately ¥31–¥48 ($0.27–$0.43) in the “RT6.25” condition (Additional file 4: Table S3, Additional file 1: Figure S6a). The “RT6.25” low-enzyme condition thus further reduced the experimental cost and slightly improved the quantitative performance.

### Quartz-Seq2 shows higher efficiency of UMI conversion and detects more biological pathways than Drop-seq

In the high-throughput single-cell RNA-seq methods, the total number of sequence reads generated by a deep sequencer is limited. Ideally, greater UMI counts should be generated from limited sequence reads because the increase in UMI count assigned to each cell leads to the detection of low-copy genes and the identification of cell-type-specific genes using statistical tests. We compared Quartz-Seq2 to the high-throughput single-cell RNA-seq method using two distinct cell types. To prepare these two distinct cell types, we cultured G6GR ES cells and Dex-treated G6GR cells because it has been reported that almost all G6GR ES cells differentiated into primitive endoderm-like cells upon dexamethasone treatment [15,23].

Drop-seq is one of the high-throughput single-cell RNA-seq methods, which can capture thousands of cells and barcoded beads in half an hour [6]. We performed Drop-seq experiments, which were validated by species-mixing analysis (Figure S11a, b). We performed Quartz-Seq2 and Drop-seq on a mixture of mouse ES cells and Dex-treated mouse ES cells [primitive endoderm (PrE) cells] and calculated the UMI conversion efficiency (Figure 3a and Additional file 1: Figure S11c). The effectiveness of Quartz-Seq2 ranged from 25% to 35% depending on the initial fastq read depth (Figure 3), which was higher than that of Drop-seq.

**Figure 3.**
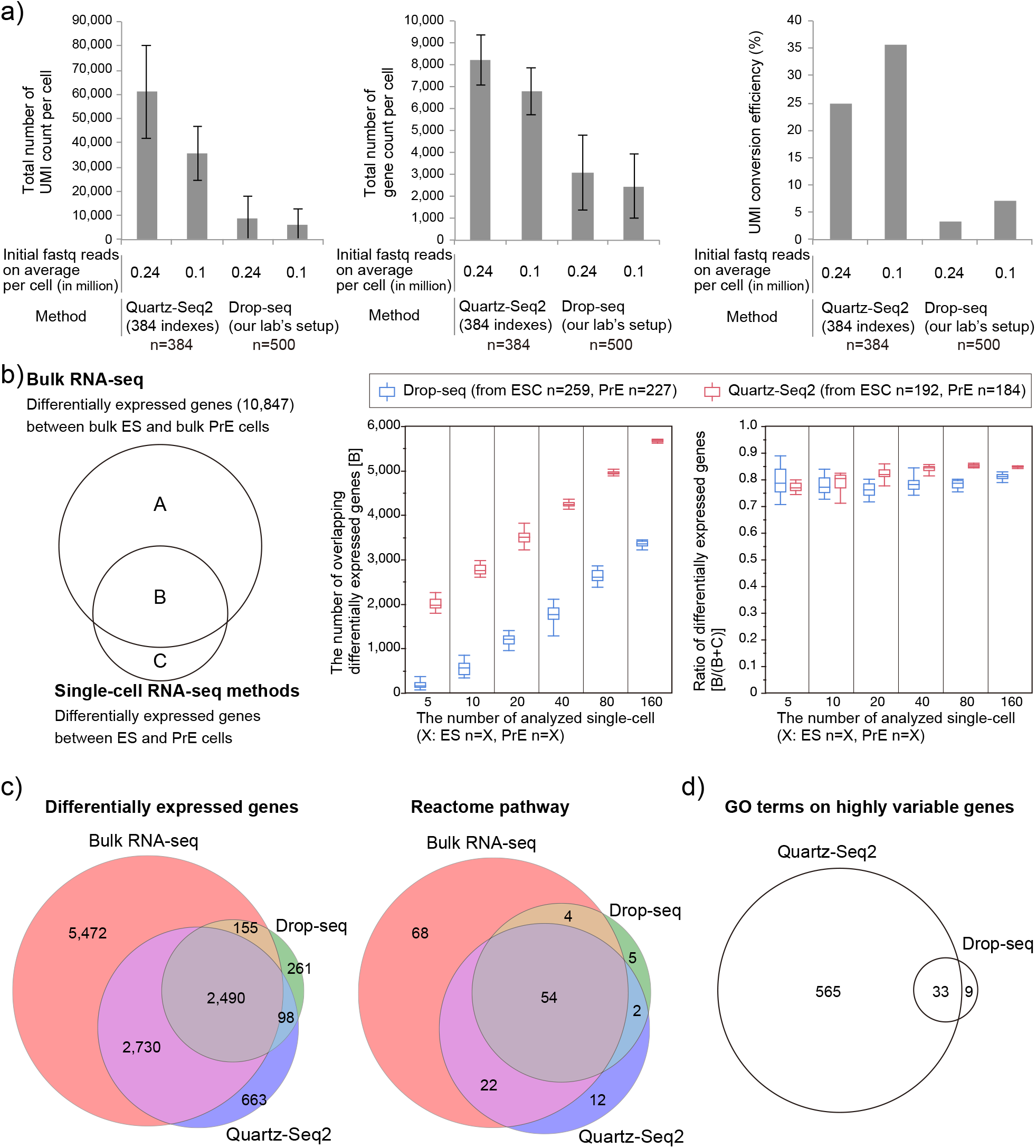
Quartz-Seq2 with high UMI conversion efficiency identified more differentially expressed genes and biological pathways. a) UMI counts, gene counts, and UMI conversion efficiency for Quartz-Seq2 and Drop-seq experiments. These values depended on the initial fastq reads on average per cell. b) We calculated overlapping differentially expressed genes between bulk RNA-seq data and single-cell RNA-seq data. We randomly picked up the indicated number of single cells and calculated differentially expressed genes 20 times. c,d) We randomly selected 100 ES cells and 100 PrE cells for each method. c) Venn diagram of genes that were differentially expressed between the ES cluster and the Dex-treated ES (PrE) cluster, as identified by Quartz-Seq2, Drop-seq, and bulk RNA-seq (left). The number of genes that differed in expression level between ES and PrE cells by at least twofold was determined (FDR < 0.05). We also present a Venn diagram of the Reactome pathway (right). d) Venn diagram of GO terms for genes with highly variable expression, as identified by Quartz-Seq2 and Drop-seq.

As Figure 3a shows, Quartz-Seq2 detected more genes with high UMI conversion efficiency. To examine the power for identifying differentially expressed genes, we performed principal component analysis (PCA) and clustering. We randomly selected single cells from ES cluster cells and PrE cluster cells. The number of genes differentially expressed between two distinct cell types was calculated (ES and PrE cells) (Additional file 1: Figure S11h). We also identified differentially expressed genes (DEGs) between bulk ES cells and that PrE cells by using conventional bulk RNA-seq. We then counted the number of overlapping DEGs between single-cell RNA-seq methods and bulk RNA-seq. We observed that this number linearly correlated with the cell number (Figure 3b). Quartz-Seq required fewer cells for the detection of overlapping DEGs. These results showed that more genes that were differentially expressed between the two cell types were identified with Quartz-Seq2 (Figure 3b, c). In addition, more biological pathways particularly associated with the differentially expressed genes were detected in Quartz-Seq2 (Figure 3c). Furthermore, we calculated the highly variable expression of genes in Quartz-Seq2 and Drop-seq, which potentially include not only genes that are differentially expressed between cell types but also genes of which the expression changes depending on the cell state in a cell type. Terms related to the cell cycle state were only enriched for the genes calculated with Quartz-Seq2 (Figure 3d). Note that simulation-based power analysis also showed that Quartz-Seq2 detected more DEGs than Drop-seq (Additional file 1: Figure S12a, see Methods). These results suggest that high UMI conversion efficiency with limited initial reads leads to more biological information being revealed, such as functional terms and biological pathways.

### Superiority of Quartz-Seq2 regarding quantitative performance in the same experimental design compared with other methods

To obtain additional evidence for the superiority of Quartz-Seq2 in terms of the UMI conversion efficiency and gene count, we compared it to other methods. Using mouse J1 ES cells, Ziegenhain et al. systematically compared the quantitative performance of several single-cell RNA sequencing (RNA-seq) methods [CEL-seq2(C1), SCRB-seq, MARS-seq, and Drop-seq] that use the UMI technique [21]. Therefore, we cultured J1 ES cells under 2i/LIF conditions in accordance with the procedure described in this previous paper [21]. We sorted J1 ES cells into five 384-well plates. Subsequently, we prepared sequence library DNA of Quartz-Seq2 (RT25) with 1,152 wells (three 384-well plates) or 768 wells (two 384-well plates) on different days. For comparison between our data and those obtained in the previous study mentioned above, we used the same analytical conditions as previously applied, such as a HiSeq sequencer platform, Read2 length, the same genome file, and the same transcript annotation file.

First, we compared the quantitative performance between Quartz-Seq2 and other methods at 0.1 million initial fastq reads on average per cell (Figure 4). The results indicate that the UMI conversion efficiency levels of Quartz-Seq2 were approximately 32.55% (Day 1) and 32.25% (Day 2); the UMI conversion efficiency of the other methods ranged from 7.11% to 22.45%. The average gene counts using Quartz-Seq2 were approximately 6,636 (Day 1) and 6,584 (Day 2), while the average gene counts for the other methods ranged from 2,738 to 5,164. We also validated the quantitative performance for external control RNA. The levels of ERCC capture efficiency for Quartz-Seq2 were approximately 6.12% (Day 1) and 6.38% (Day 2), while the ERCC capture efficiencies for the other methods ranged from 0.76% to 3.22% (Figure 4a and Additional file 1: Figure S13d). We also calculated the copy number of the detection limit of ERCC spike-in RNA. The detection limits for Quartz-Seq2 were approximately 6.82 (Day 1) and 6.57 (Day 2), while those for the other methods ranged from 13.26 to 710.24 (Additional file 1: Figure S13d).

**Figure 4.**
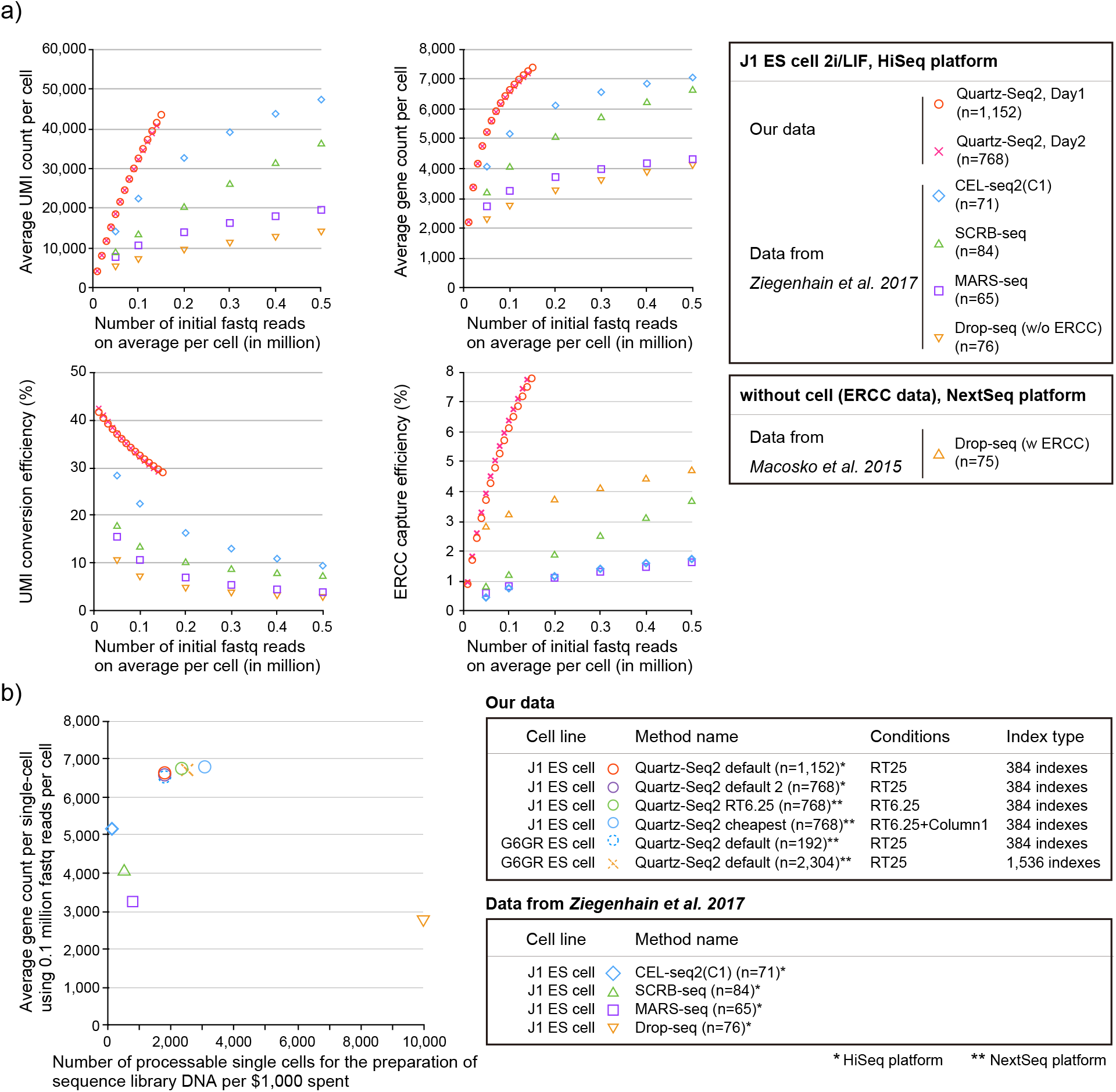
Quantitative comparison among Quartz-Seq2 and previously reported methods using embryonic stem cells. a) We determined the UMI and gene counts with Quartz-Seq2 in the “RT25” condition using J1 ES cells. We performed Quartz-Seq2 (RT25) with three sets of 384-well plates and two sets of 384-well plates on different days. We also estimated the UMI conversion efficiency of other single-cell RNA-seq methods [CEL-seq2(C1), SCRB-seq, MARS-seq, and Drop-seq] from a previous study that used mouse ES cells [21]. In our comparison, the Read2 length for transcript mapping was 45 nt for all of the methods, including Quartz-Seq2. We estimated the average UMI and gene counts and the UMI conversion efficiency with various numbers of initial fastq reads for each method. The findings indicate that, compared with the other methods, Quartz-Seq2 has a superior ability to detect UMI and gene counts from limited initial amounts of data (under 0.2 million fastq reads). b) To investigate the throughput capacity for establishing sequence library DNA, we estimated the number of processable single cells per $1,000 spent on each method: Quartz-Seq2 (384 indexes, RT25) yielded 1,785 cells, Quartz-Seq2 (384 indexes, RT6.25) yielded 2,325 cells, Quartz-Seq2 (384 indexes, RT6.25+column1) yielded 3,058 cells, Quartz-Seq2 (1,536 indexes, RT25) yielded 2,500 cells, CEL-seq2(C1) yielded 111 cells, SCRB-seq yielded 500 cells, MARS-seq yielded 769 cells, and Drop-seq yielded 10,000 cells. The UMI conversion efficiency was approximately 32.55% (n = 1,152), 32.25% (n = 768) and 32.12% (n = 192) for Quartz-Seq2 (384 indexes, RT25), 35.48% (n = 2,304) for Quartz-Seq2 (1,536 indexes, RT25), 34.04% (n = 768) for Quartz-Seq2 (384 indexes, RT6.25), 35.51% (n = 768) for Quartz-Seq2 (384 indexes, RT6.25+column1), 22.4% for CEL-seq2(C1), 13.3% for SCRB-seq, 10.6% for MARS-seq, and 7.1% for Drop-seq. The average gene count was approximately 6,636 (n = 1,152), 6,584 (n = 768) and 6,529 (n = 192) for Quartz-Seq2 (384 indexes, RT25), 6,712 (n = 2,304) for Quartz-Seq2 (1,536 indexes, RT25), 6,753 (n = 768) for Quartz-Seq2 (384 indexes, RT6.25), 6,794 (n = 768) for Quartz-Seq2 (384 indexes, RT6.25+column1), 5,164 for CEL-seq2(C1), 4,044 for SCRB-seq, 3,252 for MARS-seq, and 2,738 for Drop-seq.

We also performed simulation-based power analysis to estimate the power for identifying differentially expressed genes (Additional file 1: Figure S12b, see Methods). The true positive ratio (TPR) at the data point of 256 cells ranged from 0.93 and 0.95, and the false discovery rate (FDR) at the data point of 256 cells ranged from 0.07 to 0.09. Therefore, TPR and FDR for each method were comparable to each other. We found that Quartz-Seq2 detected more simulated DEGs than the different methods.

In particular, the UMI conversion efficiency and gene count of the Quartz-Seq2 method were significantly better than those of the other methods at approximately 0.1 million initial reads. We also estimated the UMI and gene counts and the UMI conversion efficiency at various numbers of initial fastq reads (Figure 4a). We found that Quartz-Seq2 is greatly advantageous for detecting the UMI and gene counts from limited initial amounts of data (under 0.2 million fastq reads). These results showed that the quantitative performance of Quartz-Seq2 was almost always better than that of other methods under conditions with a limited number of initial fastq reads.

### Quartz-Seq2 achieves high UMI conversion efficiency at relatively low cost

The experimental cost for sequence library preparation is an important benchmark for single-cell RNA-seq methods because it is highly correlated with the throughput of single-cell RNA-seq methods within a limited budget. In the case of using 384 indexes, the library preparation cost of Quartz-Seq2 (RT25) is ¥63 ($0.56) (Additional file 1: Figure S6a). For Quartz-Seq2 with 384 indexes, we used three purification columns per 384-well plate at the step of cDNA purification. To reduce the amount of enzyme solution in the downstream reaction after cDNA purification, we used one purification column per 384-well plate (Additional file 4: Table 3). In the previous subsection, we also described that the “RT6.25” low-enzyme condition improved the experimental cost of Quartz-Seq2. The combination of the “RT6.25” low-enzyme condition and reduction of the number of purification columns thus further improved the library preparation cost of Quartz-Seq2 (RT6.25+column1) to ¥37 ($0.32) (Additional file 1: Figure S6a). For comparison, the library preparation costs of other methods that use a cell sorter range from ¥146 ($1.30) to ¥372 ($3.29) (Figure S6a). Regarding the cost, Quartz-Seq2 is thus highly competitive with other single-cell RNA-seq methods that use a cell sorter.

To show the potential of Quartz-Seq2 for high-throughput performance, we additionally carried out Quartz-Seq2 (RT6.25+column1) and Quartz-Seq2 (RT6.25) using J1 ES cells (Figure 4b). We estimated that the number of processable single cells per 1,000 US dollars spent on Quartz-Seq2 ranges from 1,785 to 3,058 (Figure 4b). We also calculated the total cost, including the sequencing and library preparation costs, for each method. The total cost per cell is extremely low (approximately ¥71–¥131, or $0.63–$1.16) for Drop-seq. The total cost per cell is ¥97–¥183 ($0.85–$1.62) for Quartz-Seq2. Thus, the total cost of Quartz-Seq2, a cell-sorter-based method, approaches that typical of droplet-based methods (Additional file 1: Figure S6b). In summary, the Quartz-Seq2 method achieves the highest UMI conversion efficiency and high-sensitivity detection of genes under conditions with limited numbers of initial reads, providing single-cell RNA-seq data in a high-throughput manner.

### Demonstration of high-throughput Quartz-Seq2 analysis on 4,484 mouse embryonic stem cells and differentiated conditions

To demonstrate the capability of Quartz-Seq2 for quantifying the transcriptome of a large number of cells and identifying rare cell populations, we analyzed approximately 4,484 cells from a mixture of mouse ES cells and differentiation-induced cells. Cells were prestained with Hoechst 33342 and/or Calcein-AM as an indicator of DNA content and culture condition (ES or PrE cells), respectively (Figure 5a). Calcein-AM-positive and -negative cells were sorted to twelve 384-well plates (4,608 well in total) in a checkered pattern (Figure 5a). These conditions maximize the likelihood of detecting the cell doublets caused by mis-sorting. In this analysis, as an average for cells, 0.1 M initial fastq reads were effectively converted to 35,915±1,188 UMI counts in a final gene expression matrix. The UMI conversion efficiency was 35.91 ±1.18%.

**Figure 5.**
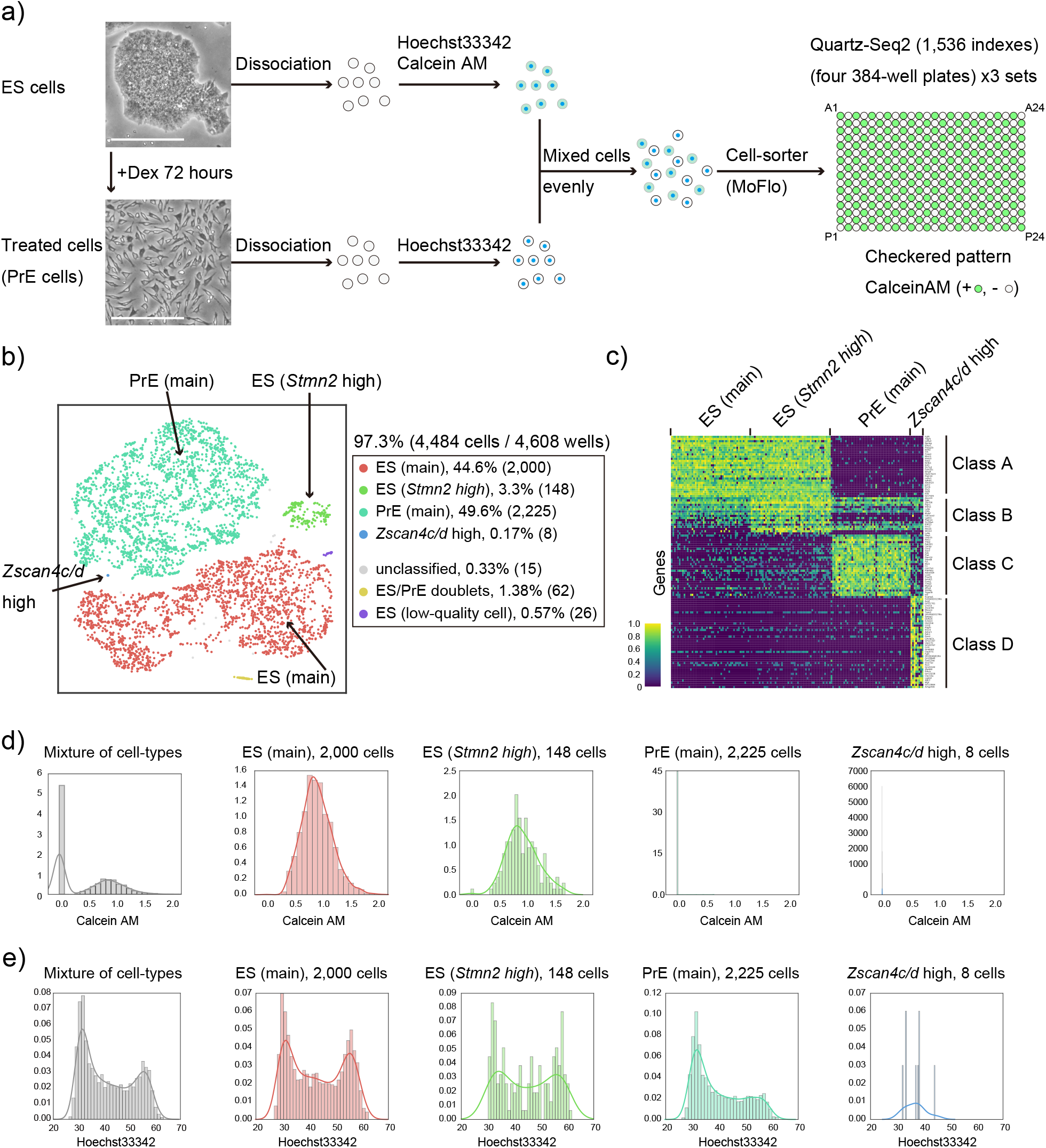
High-throughput Quartz-Seq2 analysis of 4,484 cells from mouse embryonic stem cells and differentiated cells. a) We successfully analyzed 97.3% of 4,608 wells. The procedures for cell suspension as used in this assay are shown. Cells cultured under ES-maintenance and Dex-treatment conditions were separately dissociated into single cells, stained with Hoechst 33342 and/or Calcein-AM, and mixed evenly. Calcein-AM-positive and -negative cells were sorted to 384-well plates in a checkered pattern. White scale bars represent 100 μm. b) Clustering of 4,484 single cells according to the transcriptome. Plotting of cells on t-SNE space with color labeling for each cluster. The percentage indicates the proportion of cells for each cluster relative to all cells analyzed. Numbers in parentheses indicate the numbers of cells making up the cluster. c) Marker genes for each cluster identified by Quartz-Seq2. Cluster-specific or cluster-enriched genes were calculated for each cluster, and their expression is displayed as a color in a heatmap. No more than 50 cells are shown for simplicity. d) Reconstructed distribution of Calcein-AM intensity for each cluster. The x-axis represents the intensity of Calcein-AM dye staining. e) Reconstructed distribution of Hoechst 33342 intensity for each cluster. The y-axis represents the density of cells. The x-axis represents the intensity of Hoechst 33342 dye staining.

Dimensionality reduction of the expression UMI matrix resulting from Quartz-Seq2 showed clear separation of six clusters, including the main populations of ES cells and PrE cells, as well as four small populations. We did not observe a clear batch effect among the twelve 384-well plates (Additional file 1: Figure S15). One cluster contained a relatively high proportion of mitochondrial RNA (Additional file 1: Figure S16), which was judged to reflect low-quality cells [24]. Another cluster showed high values of detected UMI counts and gene counts, and expressed both ES and PrE marker genes, which were judged to represent doublets of an ES cell and a PrE cell (Additional file 1: Figure S16). To characterize the identified populations, we determined the genes that were specific or enriched for each cluster using binomial tests (Figure 5c). Although cells in cluster 3 shared gene expression with the main population of ES cells (cluster 1), some genes including *Stmn2* and *Rhox5* were additionally expressed, suggesting that cluster 1 and cluster 3 shared the same cell type but had different cell states. Cluster 5 was a small population (0.17%, Figure 6b), but was characterized by many specific marker genes (Figure 5c), including *Zscan4c* and *Zscan4d*, which are known to be expressed in a subpopulation of mouse ES cells [25]. However, among our culture conditions, these genes were more expressed in the Dex-treatment condition than in the ES-maintenance condition. Flow cytometry information on cells in cluster 5 also showed no fluorescence of Calcein-AM, indicating that these cells were from the suspension of Dex-treated cells (Figure 5d). These observations were consistent with a previous study demonstrating that *Zscan4* was more expressed in a differentiation condition induced by the withdrawal of leukemia inhibitory factor [7].

**Figure 6.**
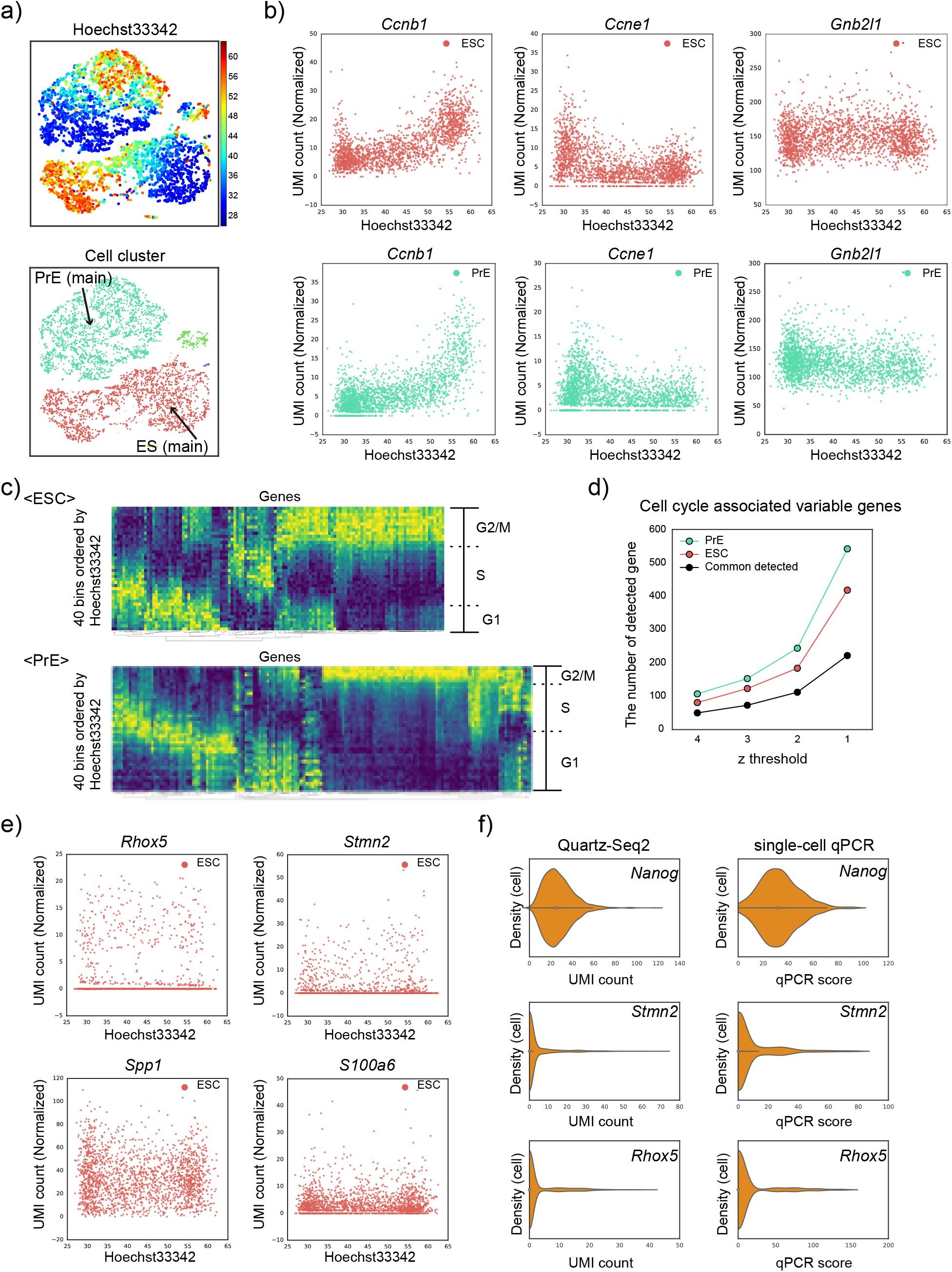
Quartz-Seq2 classified genes with variable expression within a cell type. a) Plotting cells on t-SNE space with the colors representing the intensity of Hoechst 33342 staining. b) Dependence of expression of several genes on the intensity of Hoechst 33342 staining [G_2_/M phase: *Ccnb1;* G_1_ phase: *Ccne1;* housekeeping gene: *Gnb2l1;* upper panels: ES main population (cluster 1); lower panels: PrE main population (cluster 4)]. c) Identification of cell-cycle-associated genes. Average gene expression of cells in a bin is depicted by color in the heatmap. Bins are arranged in order of Hoechst 33342 staining intensity. d) The number of cell-cycle-associated genes identified for each z-threshold. CV values from which the z-score was calculated were as follows. ES cells (z=4: 1.17, z=3: 0.91, z=2, 0.68, z=1: 0.4). PrE cells (z=4: 1.1, z=3: 0.88, z=2: 0.64, z=1: 0.39). e) Several examples for genes whose expression was variable, but not associated with the phase of the cell cycle. f) Single-cell qPCR detection of genes with variable expression.

In the sampling of cells using a cell sorter, flow cytometry information including the intensity of Hoechst 33342 staining for each cell was collected. Using this information, we reconstructed the distribution of Hoechst 33342 staining intensity for each cluster (Figure 5e). The main population of ES cells (cluster 1) showed an embryonic pattern of distribution of DNA content, which is characterized by a high ratio of cells in G_2_ and M phases to cells in the G_1_ phase. In contrast, the main population of PrE cells (cluster 4) showed a somatic pattern of distribution of DNA content. This was consistent with PrE cells being differentiated from ES cells. The cells in cluster 3, the gene expression of which was similar to that of the ES main population (cluster 1), showed an embryonic pattern rather than a somatic one. As the number of cells in cluster 5 that expressed *Zscan4c/d* was small, it was difficult to classify the observed pattern as the embryonic or somatic type. These findings indicated the usefulness of single-cell RNA-seq using flow cytometry for the reconstruction of population information after transcriptome-based clustering. For good interpretation of the distribution, a large number of cells for each cluster are required.

### Quartz-Seq2 classified genes with variable expression within a cell type

When cells were plotted on t-SNE (t-distributed stochastic neighbor embedding) space using transcriptome analysis and the intensity of Hoechst 33342 staining was depicted using a color-based scale, the gradient pattern was easily observed for clusters 1, 3, and 4 (Figure 6a), suggesting that Quartz-Seq2 was highly sensitive for detection of the cell cycle state. To examine this, we plotted the gene expression of several cell cycle markers against the intensity of Hoechst 33342 staining. We observed a strong relationship between the gene expression of several cell cycle markers and DNA content (Figure 6b). Using the intensity of Hoechst 33342 staining as reference of the cell cycle phase, we calculated the relationship between the cell cycle phase and all genes detected in Quartz-Seq2 experiments (see Methods). The results enabled us to identify numerous genes for which the expression level changed in relation to the DNA content (Figure 6c, d and Additional file 1: Figure S17). We called these genes “cell cycle associated variable genes”. These genes included cell cycle markers. Again, it was confirmed that the ratio of cells in G_1_ and S phases to those in G_2_ and M phases differed between ES and PrE cells (Figure 6c). As we showed that the efficiency of UMI conversion was high in the Quartz-Seq2 method, we examined whether 10,000 initial reads are sufficient for cell cycle analysis (Additional file 1: Figure S18). In this analysis, the final UMI count was 4,774±62 and the UMI conversion efficiency was 47.74±0.66% (n=3, 1,536 wells). Plotting cells on t-SNE space while using color to depict the intensity of Hoechst 33342 staining revealed a gradient pattern, which was positively correlated with the expression of G_2_/M-phase marker genes including *Ccnb1* and *Top2a* and negatively correlated with the expression of G_1_-phase marker genes such as *Ccne1* (Additional file 1: Figure S18). These observations demonstrated that Quartz-Seq2 with few initial sequence reads could analyze the cell cycle due to the high UMI conversion efficiency.

Within a single cell type, there are two types of cell state categories. One is the different stages of the cell cycle as mentioned above, and the other refers to cellular heterogeneity, which is only remotely related to the cell cycle phases. This means that genes with variable expression in a cell type include “cell cycle associated variable genes” and “variable genes the expression of which is less associated with the cell cycle phase” (Additional file 1: Figure S24). We determined the number of the latter group of genes by subtracting the former group from the genes with variable expression (Figure 6e). For the main population of ES cells (cluster 1), we identified tens of variable genes the expression of which is less associated with the cell cycle phase. These genes included *Spp1*, *Rhox5*, and *S100a6*, which were previously identified as genes with highly variable expression by scRNA-seq (Figure 6) [7,13,15]. We also identified *Stmn2*, *Dnmt3l*, *Tmsb4x*, and several other genes as novel variable genes the expression of which is less associated with the cell cycle phase. Single-cell RT-qPCR also showed high variability for *Stmn2* and *Rhox5* compared with that for *Nanog* (Figure 6f). In previous studies, *Sgk1* and *Actb* were identified as genes with variable expression by scRNA-seq, which analyzed cells only in the G_1_ phase or cells without measuring the intensity of Hoechst 33342 [7,15]. In this study, these were classified as cell cycle associated variable genes (Additional file 1: Figure S17).

These results show that Quartz-Seq2 globally classified genes with variable expression within a cell type into “cell cycle associated variable genes” and “variable genes the expression of which is less associated with the cell cycle phase” by using Hoechst 33342 staining intensity data obtained by flow cytometry.

### The global picture of cell-type composition in the stromal vascular fraction as revealed by Quartz-Seq2

Compared with cultured cells, fresh tissue samples tend to be composed of multiple cell types, whose sizes are highly variable. The use of fresh samples is one promising strategy to assess the robustness of high-throughput single-cell RNA-seq for various samples. Therefore, Quartz-Seq2 was applied to cells from the mouse stromal vascular fraction (SVF), which was taken from adipose tissue (Figure 7a). SVF is thought to be one of the most important sources of mesenchymal stem cells (MSCs) due to its potential for use in cell therapies and the stimulation of endogenous repair [26–29]. However, the global picture of cell-type composition in SVF has not been clarified. To achieve this, we first observed the distribution of cell size in the SVF population and found it to be broad (5–13 μm, average 6.43±1.35 μm; n=200) (Figure 7a). This was confirmed by flow cytometry data that were obtained using a cell sorter. It is known that the typical amount of total RNA in a mammalian cell is approximately 10 pg [15,17]. Thus, the average amount of total RNA in the SVF population of approximately 1.7 pg is relatively small.

**Figure 7.**
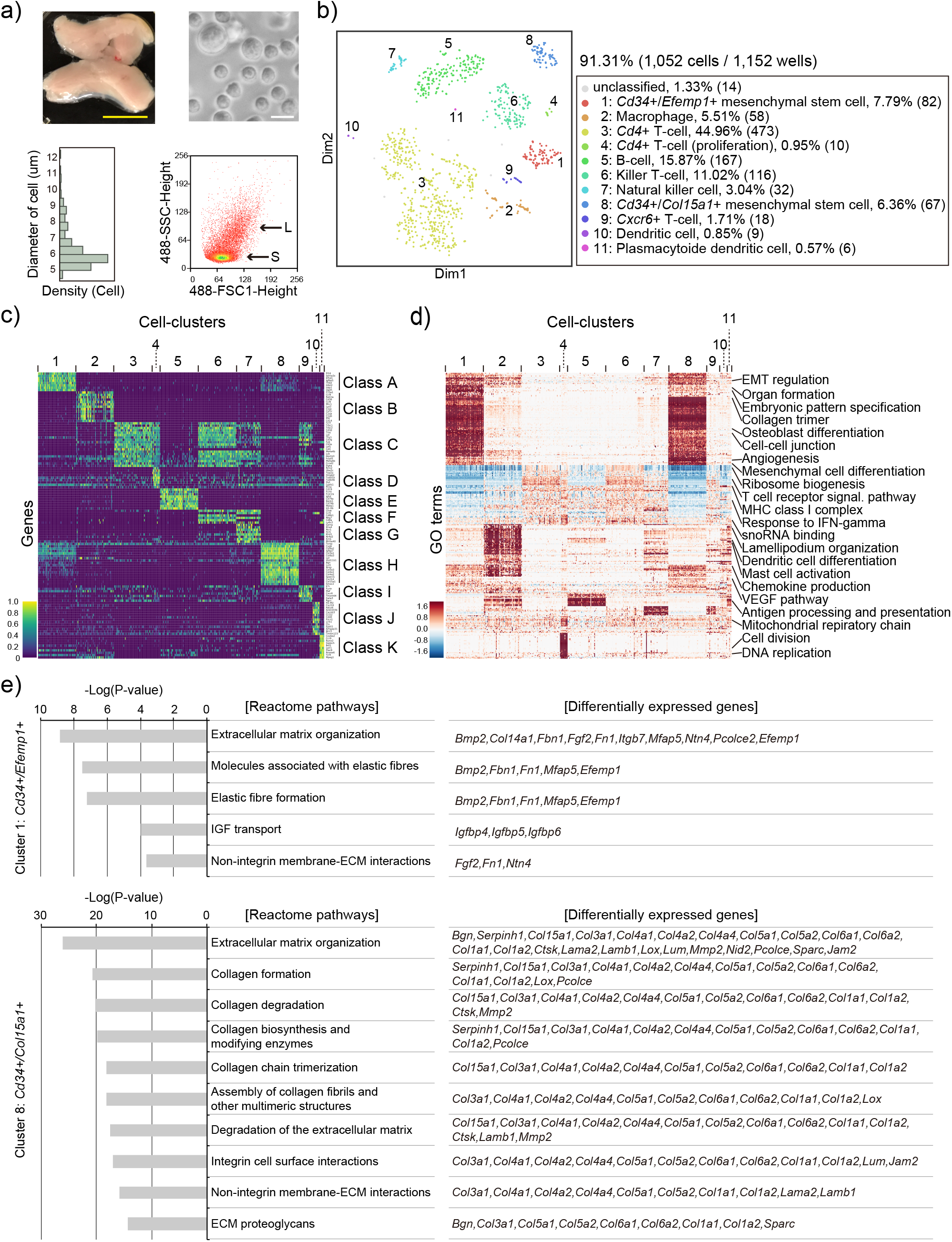
Quartz-Seq2 analysis of stromal vascular fraction (SVF) from mouse adipose tissue. a) Morphology of SVF cells. Adipose tissue from a cell suspension of SVF was prepared. Upper panels present a photograph of adipose tissues and dissociated SVF samples. Yellow scale bar represents 1 cm. White scale bar represents 10 μm. Lower panels represent the distribution of cell size information with different platforms (left: diameter of cell size using photography, right: flow cytometry information using a cell sorter). The diameter of cell size for SVF samples was 6.43±1.35 μm (n=200). b) Clustering of cells included in SVF. The transcriptome of approximately 1,000 cells was quantified by Quartz-Seq2 and clustering on t-SNE space was performed. In accordance with the genes and functional terms enriched in each cluster, the cell type was annotated. The percentage indicates the proportion of cells for each cluster relative to all cells analyzed. Numbers in parentheses indicate the numbers of cells constituting the cluster. c) Marker genes for each cluster were identified by Quartz-Seq2. Cluster-specific or cluster-enriched genes were calculated for each cluster, with their expression being displayed as color in a heatmap. No more than 50 cells are shown for simplicity. d) The results of GO-PCA analysis. Functional terms enriched in the genes with high factor loadings of PCA were calculated and the enrichment is displayed as color in the heatmap. No more than 50 cells are shown for simplicity. e) Reactome pathway with genes differentially expressed between cluster 1 and cluster 8.

We collected cells in multiple 384-well PCR plates and analyzed their transcriptome using Quartz-Seq2. In this analysis, 38,450±3513 initial fastq reads were converted to UMI counts in the final digital expression matrix. The UMI conversion efficiency was 26.85±2.70% (n=3). We observed broader distributions of UMI counts and gene counts in SVF than in ES and 10 pg of total RNA. The gene counts correlated well with side scatter (SSC) values, which were associated with cell size (Additional file 1: Figure S19).

Dimensionality reduction and clustering of single-cell transcriptome data showed the clear separation of 11 clusters (Figure 7b). To annotate the cell type for each cluster, we identified differentially expressed genes and functional terms that were either specific or enriched for each cluster (Figure 7c, d and Additional file 1: Figure S20). As Figure 7c shows, clusters 1, 2, 5, and 8–11 expressed cluster-specific genes, and clusters 3, 4, 6, 7, and 9 shared expressed genes (marker class C) as well as differentially expressed genes of classes D–G. Taking these markers and functional terms as well as previous knowledge together, each cluster was identified as follows: two types of Cd34-positive MSC (cluster 1: 7.8% of cells from SVF population, cluster 8: 6.4%), two types of Cd4-positive T cells (cluster 3: 45.0%, cluster 4: 1.0%), Cxcr6-positive T cells (cluster 9: 1.71%), B cells (cluster 5: 15.9%), killer T cells (cluster 6: 11.0%), natural killer cells (cluster 7: 3.0%), macrophages (cluster 2: 5.5%), dendritic cells (cluster 10: 0.86%), and plasmacytoid dendritic cells (cluster 11: 0.57%). We noted that the sizes of the T cells and B cells from clusters 3–5, which were measured using a cell sorter, were smaller than those of MSCs and macrophages from clusters 1, 2, and 8 (Additional file 1: Figure S19). Using immunofluorescence observation, we confirmed that the sizes of CD4-positive T cells and CD79-positive B cells were smaller in SVF (Additional file 1: Figure S19). These results show that Quartz-Seq2 with flow cytometry data is useful for defining respective cell clusters with cell size information.

Subsequently, we investigated the gene expression of receptors and ligands in cells from the SVF. The variety of receptors expressed in each cluster was similar (Additional file 1: Figure S21). However, we found a wide variety of ligands expressed in clusters 1 and 8 (Additional file 1: Figure S21). It is known that MSCs secrete many paracrine mediators, which have a therapeutic effect [30,31]. These results suggest that clusters 1 and 8 are MSCs, which secrete many paracrine mediators.

We focused on these two potential MSC clusters (1 and 8) because the heterogeneity of the MSC population has been discussed in many papers [32,33]. By using Quartz-Seq2, clusters 1 and 8 were shown to have the *Cd31−*/*Cd34*+/*Cd45−* phenotype (Additional file 1: Figure S22). In a previous study, qPCR-based single-cell transcript analysis of the *Cd31−*/*Cd34*+/*Cd45−* population of SVF was reported. By analyzing 140 genes, the authors identified the *Cd55*+/*Dpp4*+/*Cd31*−/*Cd34*+/*Cd45−* cell population and showed that the administration of this population was effective for normalizing diabetic wound healing in mice [32]. They identified 13 marker genes for the *Cd55*+/*Dpp4*+/*Cd31*−/*Cd34*+/*Cd45*− population in mouse SVF. We compared the expression of those 13 genes between clusters 1 and 8 (Additional file 1: Figure S23). The results showed that almost all of the genes were more highly expressed in cluster 1 than in cluster 8, suggesting that the cluster 1 population was similar to the *Cd55*+/*Dpp4*+/*Cd31−*/*Cd34*+/*Cd45−* population. To further analyze the relationship between our identified clusters and previously reported heterogeneity of the MSC population, we examined the expression of typical MSC markers (*CD90*, *CD105*, *PDGFRa*, and *Sca-1*) [32,34] as well as subpopulation-specific markers (*Pou5f1*, *Nanog*, *Sox2*, *Tnnt2*, and *Myog*) [35,36]. *PDGFRa* (*Pdgfra*) and *Sca1* were expressed in both clusters 1 and 8, whereas *Cd90* was expressed at a higher level in cluster 8 than in cluster 1, and *Cd105* was not strongly expressed in either cluster. *Cd90* and *Cd105* were, however, not specific to these MSC clusters. Neither pluripotent markers (*Pou5f1*, *Nanog*, and *Sox2*) nor skeletal and cardiomyogenic markers (*Tnnt2* and *Myog*) were detected in either cluster (Additional file 1: Figure S22). Collectively, our transcriptome analysis showed that the MSC population is divided into two clusters, suggesting that there is less heterogeneity of MSCs in the SVF than expected.

Next, we investigated the difference between the two MSC clusters in SVF. We detected 182 differentially expressed genes between clusters 1 and 8 (Additional file 5: Table S4). Cluster 8 was characterized by the enriched expression of genes that encode extracellular matrix proteins, including collagens, *Hspg2*, and *Bgn*. By contrast, cluster 1 was characterized by the expression of *IGFBPs* (*Igfbp4*, *Igfbp5*, *Igfbp6*), prostaglandin endoperoxidase synthases (*Ptgs2*), and secreted factors *Fgf2* and Bmp2. Notably, the expression of stem-cell-characteristic genes *Aldh1a3*, *Fgf2*, *Bmp2* and *Tgfbr2* were identified as important modules in cluster 1, suggesting that this cluster included more stem-like phenotypes that secrete medicinal factors.

To leverage the power of the entire transcriptome analysis, we performed an enrichment analysis of biological pathways using the 182 genes differentially expressed between the two clusters (Figure 7e, Additional file 6: Table S5). Cluster 1 was characterized by elastic fiber pathways and IGF transport pathways. Elastic fibers are constituents of various connective tissues and are related to vascular function [37]. Growth factors including FGF2, BMP2, IGFBPs, and PGE2 regulate growth and differentiation [38]. A tightly linked network of collagen metabolism pathways was found to be highly enriched in cluster 8. These results suggest that two MSC clusters of SVF have different functions. Cluster 1 could feature medicinal cells characterized by the expression of growth and development-related genes. In contrast, cluster 8 may be characterized by the production of extracellular matrix.

Taking the obtained findings together, our transcriptome analysis by Quartz-Seq2 successfully identified cell populations that consisted of various types of both large and small cells, which were isolated from *in vivo* tissue, and demonstrated that fresh SVF contains two closely related types of MSCs that have distinct characteristics.

## Discussion

The rate of conversion from initial reads to UMI counts was low, especially in high-throughput single-cell RNA-seq. Ideally, greater UMI counts should be generated from limited sequence reads because the increase in UMI count assigned to each cell leads to the detection of low-copy genes and the identification of cell-type-specific genes using statistical tests. To overcome these issues, we developed a novel high-throughput single-cell RNA-seq method, Quartz-Seq2. We calculated the UMI conversion efficiency, which indicated how effectively initial reads can be converted to UMI counts. The UMI conversion efficiency of Quartz-Seq2 was 1.44–4.55-fold higher than those of other single-cell RNA-seq methods (Figure 4). Quartz-Seq2 could detect 1.28–2.41-fold more genes from limited fastq reads and for a lower cost (Figure 4). Moreover, we decreased the cDNA preparation cost per cell of Quartz-Seq2 by 97.5%–98.5% compared with that of Quartz-Seq (Figure S6a). The total cost (cDNA preparation cost and sequence cost per cell) of Quartz-Seq2 as a cell-sorter-based method approached that of a droplet-based method (Additional file 1: Figure S6b). Furthermore, the use of evaporation-preventing oil might reduce the cDNA preparation cost of Quartz-Seq2 by 75% [2]. By cell barcoding, Quartz-Seq2 can pool cDNA of up to 1,536 individual cells into one mixture. Streamlined and simplified experimental processes of Quartz-Seq2 allowed us to handle thousands of single cells within a few days. Quartz-Seq2 also showed high accuracy and high reproducibility, leading to precise measurement of the transcriptome from single cells (Figure 2d and e). We demonstrated Quartz-Seq2 analyses on a total of approximately 9,000 mouse ES cells as an *in vitro* sample, and approximately 1,000 cells from mouse SVF samples including MSCs as an *in vivo* sample.

We proposed that UMI conversion efficiency could be a useful variable for evaluating performance for the further development of high-throughput single-cell RNA-seq methods with shallow reads. The increase of initial sequence reads by additional sequencing does not cancel out the low efficiency of UMI conversion at an equal rate. For example, approximately 24,000 fastq reads were converted to approximately 11,000 UMI counts in Quartz-Seq2, while Drop-seq required approximately 169,000 fastq reads to generate the same UMI counts. Under these conditions, the difference of UMI conversion efficiency was 2.5-fold but the difference of required fastq reads to generate the same UMI count was 7-fold. This is because UMI counts do not increase linearly with an increase of initial sequence reads due to UMI filtering. Unfortunately, the rate of increase of sequence throughput was lower than that of the processing ability for single cells. It will thus continue to be important to utilize limited initial fastq reads for high-throughput single-cell RNA-seq methods.

Quartz-Seq2 is based on poly-A tagging, which is one of the strategies of converting first-strand cDNA to amplified cDNA [19,39]. However, the poly-A tagging efficiency itself has not been improved for single-cell RNA-seq. In this study, three molecular biological improvements of Quartz-Seq2 contributed to the increase of amplified cDNA, which led to high UMI conversion efficiency (Figure 2 and Additional file 1: Figure S1). In addition, the improvement of poly-A tagging steps was the most efficient approach to increase amplified cDNA for Quartz-Seq2 (up 360%, Additional file 1: Figure S1). Several previous studies improved the efficiency of conversion from an mRNA molecule to amplifiable cDNA, and showed that the increase of amplified cDNA is a good guidepost to improve UMI counts or quantitative performance for single-cell RNA-seq [3,14,15]. These results suggest that improved poly-A tagging was the most important feature for the high UMI conversion efficiency of Quartz-Seq2.

Several improvements proposed in this study could contribute to the further development of other single-cell RNA-seq methods, as follows. (1) We showed that a decrease of enzyme concentration in RT solution led to decreases in technical error and cDNA preparation cost. (2) We applied the Sequence–Levenshtein distance for the design of cell barcodes containing RT primer sets. The use of primer sets designed in this study allows the user to correct mutations of at most two nucleotides of a cell barcode, including substitution, insertion, or deletion; the correction capability of these primer sets is higher than that of barcode sets used in previous studies [1,3]. Such correction of the cell barcode increased the average UMI count by 3%-5%. (3) We hope that the spin-down collection system developed in this study can be applied to other cell-sorter-based single-cell RNA-seq methods [2,3,11,12]. (4) In Quartz-Seq2, we improved the efficiency of poly-A tagging itself (Figure 2). Several single-cell RNA-seq methods including Quartz-Seq were based on poly-A tagging [4,5,10,15,19,40]. Such improvement may directly contribute to these methods having increased quantitative performance.

We achieved a marked improvement in UMI conversion efficiency at a low cost from initial reads to UMI counts, allowing us to analyze 3-to 10-fold more single cells in limited sequence experiments. However, it is difficult to sort more than 20,000 single cells in a day when using one cell sorter. While there is greater scalability for cell sampling in droplet-based single-cell RNA-seq methods than in cell-sorter-based methods, these latter methods can utilize additional information obtained in flow cytometry [41,42], which cannot be determined from transcriptome data alone. These two types of method provide complementary approaches for investigating complex biological phenomena. C1 Single-Cell Auto Prep System (Fluidigm) is another widely used platform for single-cell RNA-seq methods [43]. CEL-seq2 shows high UMI conversion efficiency, so it is a very convenient method for users of the C1 platform as a single-cell RNA-seq method based on UMI count [3]. Quartz-Seq2 cannot be performed with the C1 platform, but it can assimilate 1,536 cell barcodes in a cell sorter in a high-throughput manner. We think that both methods can be effective as long as the most appropriate one is selected for each situation.

In this study, we showed that Quartz-Seq2 has advantages in gene detection and the identification of biological pathways via high UMI conversion efficiency (Figure 3). Moreover, we analyzed thousands of single cells from all cell cycle phases within a cell type (Figure 6). These specifications with Quartz-Seq2 allow us to perform the global classification of genes with variable expression in a cell type into “cell cycle associated variable genes” and “variable genes the expression of which is less associated with the cell cycle phase”.

While Hoechst 33342 is used as an indicator of DNA content, FUCCI reporter systems monitor the ratio of activities for two different cell-cycle-associated proteins. Therefore, the latter has an advantage in cell cycle analysis via providing more detailed resolution, although the introduction of reporter constructs into the cells cannot be applied for all purposes, especially for the analysis of human samples. A previous study utilizing the FUCCI fluorescent reporter system reported that competence to respond to specific differentiation signaling was limited to only an early or a late window of the G_1_ phase in human ES cells [44]. In the future, combining Quartz-Seq2 and FUCCI/FUCCI2 or other fluorescent reporter systems should lead to an understanding of the global picture of differentiation dynamics regarding competence, response, transition, and commitment because Quartz-Seq2 can analyze changes of cell state for thousands of cells.

In summary, Quartz-Seq2 can be used to obtain continuous data on cell states because of the large number of cells with which it deals and the high efficiency of use of initial sequence reads. Quartz-Seq2 can facilitate investigation of the cell state within a cell type, such as gradated or stochastic changes of the cell population in organism development and disease progression.

## Conclusions

In this study, we developed a high-throughput single-cell RNA-seq method, Quartz-Seq2, which can analyze cells numbering up to 1,536 that are pooled together in a single sample. Quartz-Seq2 allows us to effectively utilize initial sequence reads from a next-generation sequencer. The UMI conversion efficiency in Quartz-Seq2 ranged from 32% to 35%, which is much higher than for other single-cell RNA-seq methods (7%–22%). This was caused by the improvements in several molecular biological steps including poly-A tagging. The technical gene expression variability of Quartz-Seq2 was close to the theoretical variability of a Poisson distribution. As we showed in the analysis of SVF and the ES/PrE mixture, cell types in the population were identified with marker genes and functional terms that characterized each cell type. We identified two types of Cd34-positive MSCs in SVF, namely, those that express numerous transcription-factor-encoding and secreted-protein-encoding genes specific to each. Quartz-Seq2 can also be used to provide continuous data on cell states because of the large number of cells with which it deals and the high efficiency in the use of initial sequence reads. Quartz-Seq2 should facilitate investigations of the cell state within a cell type, such as gradated or stochastic changes of cell populations in organism development and disease progression.

## Methods

### Cell culture

Mouse embryonic stem cells were cultured as described previously [15]. Briefly, 1.0 x 10^5^ cells were seeded on a 60-mm dish coated with gelatin (0.1%). Cells were maintained in GMEM-based medium containing 10% FBS and 1000 units/ml leukemia inhibitory factor (Millipore ESGRO). The cell line used in this study was G6GRGFP, which was established in a previous study [23]. It has been reported that almost all G6GRGFP differentiated into primitive endoderm-like cells upon dexamethasone treatment [15,23]. To differentiate ES cells into primitive endoderm-like cells, 1.0 x 10^5^ cells were seeded on a 60-mm dish coated with gelatin and cultured in GMEM-based medium containing 10% FBS, 1000 units/ml LIF, and 100 mM dexamethasone for 72 h. We confirmed that almost all of the Dex-treated G6GR ES cells differentiated into primitive endoderm-like cells. Moreover, we cultured J1 ES cells in accordance with a procedure described in a previous report [21].

### Cell staining

To identify dead or damaged cells, propidium iodide (PI) was added to the cell suspension (final concentration 1–2 μg/ml). As an indicator of an undifferentiated state in a suspension containing a mixture of cells, Calcein-AM was used. Suspensions for cells cultured under maintenance and differentiation conditions were prepared independently, and cells in the maintenance condition were treated with 1 μg/ml Calcein-AM for 10 min on ice. After washing with PBS, cell suspensions of ES cells and PrE cells at the same concentration were mixed. Hoechst 33342 was used as an indicator of the DNA content in a cell. The procedures were performed as described previously [15].

### Single-cell preparation for stromal vascular fraction

These experiments were carried out in accordance with the protocol approved by the National Institute of Advanced Industrial Science and Technology (AIST) Animal Care and Use Committee and the RIKEN Animal Experiment Committee. Subcutaneous fat tissues from three-to four-month-old ICR male mice (n=3 per sample, two biological replicate samples) were minced into small pieces and incubated with 0.4 U/mL collagenase NB4G (Serva) at 37°C for 35 min in a shaking water bath. The digested solution was sequentially filtered through 100-μm and 40-μm cell strainers (Corning), followed by centrifugation at 250 × *g* for 5 min to remove mature adipocytes. The pellet was treated with erythrocyte lysis buffer (BD Biosciences) and centrifuged at 180 × *g* for 5 min. The nucleated cells were suspended in HBSS with 0.1% BSA, filtered through a 20-μm cell strainer (pluriSelect), and then kept on ice (Cell solution A). The cell aggregates that did not pass through the 20-μm strainer were further treated with Accutase (Thermo Fisher Scientific) at 37°C for 15 min to dissociate them into single cells, centrifuged at 180 × *g* for 5 min, and suspended in HBSS with 0.1% BSA (Cell solution B). Cell solutions A and B were mixed and again filtered through the 20-μm cell strainer, followed by centrifugation at 180 × *g* for 5 min. The pellet was resuspended with HBSS with 0.1% BSA, stained with PI, and used for single-cell analysis.

### RNA preparation

Total RNA was purified from cultured cells using Direct-zol RNA MiniPrep kit (Zymo Research) with TRIzol RNA Isolation Reagents (Thermo). We measured the concentration of purified total RNA using a NanoDrop 1000 Spectrophotometer (Thermo). We confirmed that the RNA integrity number of total RNA was over 9.5 using an Agilent RNA 6000 Nano Kit (Agilent). Estimation of the average amount of total RNA per single cell for respective samples was performed in accordance with our previous study [15]. For high-throughput single-cell RNA-seq, we prepared total RNA from a single cell with ERCC spike-in RNA. First, we diluted the ERCC spike-in RNA tenfold. We then added 6 μL of 1:10 diluted ERCC spike-in RNA per 10 μg of total RNA. We used diluted 10 pg of total RNA with ERCC spike-in RNA for the technical validation of Quartz-Seq2. For single cells, we used the same concentration of ERCC spike-in RNA.

### Bulk RNA-seq methods for populations of cells

We prepared sequence library DNA with 1 μg of total RNA using NEBNext Poly(A) mRNA Magnetic Isolation Module and NEBNext Ultra Directional RNA Library Prep Kit. The total RNA did not contain ERCC spin-in mix I. In addition, we used SuperScript III instead of ProtoScript in the reverse-transcription step and KAPA HiFi DNA polymerase instead of NEBNext High-Fidelity PCR DNA polymerase in the PCR step. The resulting sequence library DNA was analyzed by HiSeq2500.

### Design of cell barcodes

The selection of 384 or 1,536 sequences for v3.1 and v3.2 barcode primer sets was performed as described below. First, 1,582 or 4,714 candidate sequences were created using the DNABarcodes package of R Bioconductor for the v3.1 set with 14-mer and the v3.2 set with 15-mer, respectively (version: 1.0.0). To reduce the loss of reads converted into UMI counts, we applied the Sequence–Levenshtein distance as an edit distance in order to maximize the ability to correct errors that occur during the synthesis of oligonucleotides or sequencing [22]. The minimum distance between any two sequences was controlled to 5, leading to the correction of a maximum of two errors of substitution, insertion, or deletion. As the base composition of the created sequences was not uniform, we selected 384 or 1,536 sequences from among 1,582 or 4,714 created sequences so that the variance of base composition decreased. These sequences are listed in Additional file 3: Table S2.

### Single-cell collection using flow cytometry

Stained cells were analyzed using flow cytometry SH800 (Sony) or MoFlo Astrios EQ (Beckman Coulter). For SH800, we used 130-μm microfluidic sorting chips. For MoFlo Astrios EQ, we used 100-μm nozzle sizes. Each cell sorter was equipped with a custom-made splash-guard to prevent contamination from unexpected droplets between the target well and neighboring wells. We used two sets of RT primer for Quartz-Seq2. The set of v3.1 RT primers has 384 kinds of unique cell barcodes, with a length of 14 nucleotides (OPC purification, FASMAC). The set of v3.2 RT primers has 1,536 kinds of unique cell barcodes, with a length of 15 nucleotides (OPC purification, Sigma). The set of v3.1 RT primers corresponds to one set of the 384-well PCR plate with lysis buffer, the wells of which have unique barcodes. The set of v3.2 RT primers corresponds to four sets of the 384-well PCR plate with lysis buffer. The RT primer position in the 384-well plate and the sequence were as described in Additional file 3: Table S2. Single cells were isolated in the 384-well PCR plate with 1 μL of lysis buffer (0.1111 μM respective RT primers, 0.12 mM dNTP mix, 0.3% NP-40, 1 unit/μL RNasin plus) containing ERCC spike-in RNA. During single-cell sorting, the 384-well PCR plate was kept on a 384 aluminum stand at 4°C. For Moflo Astrios EQ, we used G5498B-060 Insert 384 Eppendorf twin.tec PCR as a 384 aluminum stand (Agilent). For SH800, we used an SH800 384 aluminum stand (Sony). Immediately after the cell collection, the plate was temporarily sealed with LightCycler 480 Sealing Foil (Roche) and the sealed 384-well PCR plate was centrifuged at 10,000 g and 4°C for 1 min using TOMY MX307, equipped with a Rack-in-Rotor and centrifugation rack PCR96–02. These steps were very important to collect the droplet with a single cell in lysis buffer in an efficient manner. By altering the volume of lysis buffer from 0.4 μL (as described in the original Quartz-Seq paper) to 1 μL (Quartz-Seq2), bubbling of lysis buffer was not required before single-cell sorting and the lysis buffer could easily be handled. We then peeled open the temporary seal and re-sealed it with Agilent PlateLoc Thermal Microplate Sealer (Agilent). We agitated the plate at 2,600 rpm and 4°C for 1 min using ThermoMixer C (Eppendorf), after which we centrifuged the plate again. The resulting 384-well plate was then immediately cryopreserved at −80°C and maintained under such conditions until subsequent reverse transcription for cell barcoding. After cryopreservation, we performed subsequent whole-transcript amplification using the cryopreserved 384-well PCR plate within a few months.

### Whole-transcript amplification of Quartz-Seq2

Cryopreserved 384-well plates with single-cell lysate were centrifuged at 10,000 g and 4°C for 1 min. Subsequently, we denatured total RNA in each 384 plate at 70°C for 90 s and hybridized the RT primer to poly-adenylated RNA at 35°C for 15 s using the C1000/S1000 thermal cycler. The resulting plates were again centrifuged at 10,000 g and 4°C for 1 min. Next, the plates were placed on the 384 aluminum plate at 0°C. We peeled away the seal and added 1 μL of RT premix (2x Thermopol buffer, 5 units/μL SuperScript III, 0.55 units/μL RNasin plus) to 1 μL of lysis buffer for each well using a Mantis microfluidic dispensing system (Formulatrix) or a 384 Transfer Plate system (1859–384S, Watson). The above RT solution was used for the RT25 condition. For the RT100 condition, we used the following RT solution: 2x Thermopol buffer, 20 units/μL SuperScript III, and 2.2 units/μL RNasin plus. We sealed the plates again and agitated them at 2,600 rpm and 4°C for 1 min. The plates were then centrifuged at 10,000 g and 4°C for 1 min. We then performed reverse transcription at 35°C for 5 min and 50°C for 50 min. The reverse transcription was stopped at 70°C for 15 min. Then, the plates were placed on a prechilled aluminum block, after which we peeled off their seals. Subsequently, we turned the plates upside down on the assembled collector type A or type B (supplemental figure). We mainly used type A. We centrifuged the plates with an assemble collector at 3,010 g and 4°C for 3 min with swing-bucket rotors. Subsequently, we collected the cDNA solution into a disposable reservoir. Typically, we obtained 650–700 μL of cDNA solution from one 384-well PCR plate. We purified and concentrated the cDNA solution using the DNA Clean & Concentrator™-5 kit (Zymo Research). We used three purification columns for one 384-well PCR plate in the case of the v3.1 RT primer system (384-cell barcode). Purified cDNA was extracted into 20 μL of nuclease-free water from one column purification and transferred into an eight-linked PCR tube (TaKaRa). The PCR tubes were placed on an aluminum PCR stand at 0°C. We added 25 μL of TdT solution [1x Thermopol buffer, 2.4 mM dATP, 0.0384 units/μL RNase H (Invitrogen), 26.88 units/μL terminal transferase (Roche)] into 20 μL of extracted cDNA using a pipette at 0°C. The resulting 45 μL of TdT solution was mixed with a pipette at 0°C or ThermoMixer at 2,000 g and 0°C for 1 min. Immediately thereafter, the PCR tubes were centrifuged at 10,000 g and 0°C for 1 min. We used a C1000/S1000 thermal cycler equipped with the 96-Deep Well Reaction Module for the following steps. The PCR tubes were placed on the block of the thermal cycler, which had been prechilled to 0°C. We then performed a poly-A tailing reaction at 37°C for 75 s. The solution was inactivated at 65°C for 10 min. The PCR tubes were placed on an aluminum PCR stand at 0°C. We then dispensed approximately 11 μL of solution into four wells from 45 μL of TdT solution. We added 46.16 μL of PCR I premix (1.08492x MightyAmp Buffer version 2, 0.06932 μM Tagging primer, 0.05415 units/μL MightyAmp DNA polymerase) to 11 μL of TdT solution for the respective wells of the PCR tube. MightyAmp DNA polymerase, which used in Quartz-Seq and Quartz-Seq2, are marketed as Terra PCR Direct polymerase [15]. We performed gentle inversion mixing on the resulting solution in the PCR tube. The tubes were then centrifuged at 10,000 g and 4°C for 1 min. Subsequently, the solution was mixed with ThermoMixer at 2,000 rpm and 4°C for 2 min. Then, we spun down the tube again. Next, we denatured the solution at 98°C for 130 s and hybridized Tagging primer to poly-A-tailed cDNA at 40°C for 1 min. After that, we performed the “Increment step” by heating to 68°C at 0.2°C every second and performed second-strand synthesis at 68°C for 5 min. The tubes were placed on an aluminum PCR stand at 0°C. We added 50.232 μL of PCR II premix (0.99697x MightyAmp Buffer version.2, 1.8952 μM gM primer) to 56.16 μL of PCR I solution. We performed gentle inversion mixing on the resulting solution in the PCR tube. The tubes were then centrifuged at 10,000 g and 4°C for 1 min. Subsequently, the solution was mixed with ThermoMixer at 2,000 rpm and 4°C for 2 min, after which we spun down the tube again. We then placed it on the block of the thermal cycler at 68°C. Subsequently, we amplified the cDNA for 11 cycles under the following conditions: 98°C for 10 s, 65°C for 15 s, and 68°C 5 min. We then incubated the tube at 68°C for an additional 5 min. Finally, we transferred all of the PCR solution, derived from one 384-well PCR plate, to a 50-mL Polypropylene Centrifuge Tube (Watson). Typically, we obtained approximately 1.2 mL of PCR solution per 384-well PCR plate. We added 32 μL of 3 M sodium acetate (pH 5.2) and 6420 μL PB-Buffer (Qiagen) to the PCR solution. The mixture was then purified using a MinElute Spin Column (Qiagen). Purified cDNA was extracted into 40 μL of nuclease-free water. We additionally purified the cDNA with 32 μL of Ampure XP beads. Finally, we obtained 32 μL of purified cDNA. We checked the length distribution of amplified cDNA with an Agilent High Sensitivity DNA Kit (Agilent). The typical average size of the amplified cDNA in Quartz-Seq2 was approximately 1,400 bp (Additional file 1: Figure S2c). Primer sequences are listed in Additional file 2: Table S1.

In the case of the usage of the v3.2 RT primer in Quartz-Seq2, we modified the above steps as follows. After reverse transcription, we collected cDNA solution into a disposable reservoir from four sets of 384-well plates, which corresponded to 1,536 wells. We purified and concentrated the cDNA solution using eight purification columns for four 384-well PCR plates in the case of the v3.2 RT primer system. In the PCR step, we amplified cDNA for nine cycles with the following conditions: 98°C for 10 s, 65°C for 15 s, and 68°C for 5 min. Finally, we transferred all of the PCR solution derived from four 384-well plates to a 50-mL Polypropylene Centrifuge Tube (Watson). Typically, we obtained approximately 3.5 mL of PCR solution per four 384-well PCR plates. We added 88 μL of 3 M sodium acetate (pH 5.2) and 17.6 mL of PB-Buffer (Qiagen) to the PCR solution. The mixture was purified using the MinElute Spin Column (Qiagen). Subsequently, cDNA was again purified using Ampure XP magnetic beads.

For the Quartz-Seq1-like reaction, we performed the following procedure in accordance with our previous study. We added 1 μL of RT premix (2x PCR buffer, 5 units/μL SuperScript III, 0.55 units/μL RNasin plus) to 1 μL of lysis buffer. We then performed reverse transcription at 35°C for 5 min and 45°C for 20 min. This reverse transcription was stopped at 70°C for 15 min. We added 5 μL of ExoIB solution (1.6x Exonuclease I buffer, 3.2x PCR buffer, 16 mM DTT) and 20 μL of TdT solution [1x PCR buffer, 3 mM dATP, 0.0384 units/μL RNase H (Invitrogen), 33.6 units/μL terminal transferase (Roche)] into 20 μL of extracted cDNA using a pipette at 0°C. The PCR tubes were placed on the block of the thermal cycler that had been prechilled to 0°C. We performed a poly-A tailing reaction at 37°C for 75 s. We then denatured the solution at 98°C for 130 s and hybridized Tagging primer to poly-A-tailed cDNA at 40°C for 1 min. After that, we performed second-strand synthesis at 68°C for 5 min. Subsequently, we amplified the cDNA via a PCR reaction for 12 cycles.

### Preparation of truncated sequence adaptor

The truncated sequence adaptor was composed of an rYshapeP5 primer (HPLC-purified) and rYshapeP7LT primers (HPLC-purified), which had a TruSeqLT-compatible pool barcode. We prepared 100 μM respective primers with adaptor buffer (10 mM Tris-HCl pH7.8, 0.1 mM EDTA pH8.0, 50 mM NaCl). We added 5 μL of 100 μM rYshapeP5 primer and rYshapeP7LTxx primer into a single PCR tube. We denatured the solution at 90°C for 90 s. After that, we achieved annealing by cooling to 10°C by 0.5°C every 30 s and then maintaining the sample at 4°C. We then placed the tube on an aluminum PCR stand at 0°C. Subsequently, we added adaptor buffer, which was prechilled at 0°C, to 10 μL of 50 μM truncated adaptor. Finally, we obtained about 50 μL of 10 μM truncated adaptor. We cryopreserved 1 μL of 10 μM truncated adaptor in a PCR tube at −80°C until usage in the adaptor ligation step. Primer sequences are listed in Additional file 2: Table S1.

### Sequence library preparation of Quartz-Seq2

We added 130 μL of nuclease-free water including 5–10 ng of amplified cDNA into a Crimp-Cap microTUBE with AFA Fiber. We then performed cDNA fragmentation using an LE220 Focused-ultrasonicator (Covaris) under the following conditions: duty factor 15%, peak incident power 450 W, cycles per burst 200, and treatment time 80 s. We purified and concentrated the cDNA solution using the DNA Clean & Concentrator™-5 kit. Purified cDNA was extracted into 10 μL of nuclease-free water from one column purification and transferred into an eight-linked PCR tube (TaKaRa).

We then added 2 μL of End-Repair premix [1.4 μL of End repair & A-tailing buffer and 0.6 μL of End repair & A-tailing Enzyme (KAPA Biosystems)] to 10 μL of fragmented cDNA solution. Subsequently, we mixed the solution by pipetting on an aluminum PCR stand at 0°C. The PCR tubes were placed on the block of the thermal cycler, which had been prechilled to 20°C. We then incubated the tubes at 37°C for 60 min and 65°C for 30 min. After this, we added 2 μL of adaptor buffer (1.5 μM truncated adaptor, 10 mM Tris-HCl pH7.8, 0.1 mM EDTA pH8.0, 50 mM NaCl) and 8 μL of ligation premix [6 μL of ligation buffer, 2 μL of DNA ligase (KAPA Biosystems)] at 4°C. The 22 μL of ligation solution was well mixed by pipetting at 4°C. Then, we performed adaptor ligation at 20°C for 15 min. After ligation, we added 18 μL of Ampure XP beads to 22 μL of adaptor ligation solution and mixed them well. Adaptor-ligated cDNA was extracted into 20 μL of nuclease-free water. We added 32 μL of PCR premix [25 μL of 2xKAPA HiFi ready mix, 1.75 μL of 10 μM TPC2 primer (HPLC-purified), 10 μM P5-gMac hybrid primer (HPLC-purified)] to 18 μL of adaptor-ligated cDNA. We denatured the solution at 98°C for 45 s. Subsequently, we amplified cDNA for eight cycles under the following conditions: 98°C for 15 s, 60°C for 30 s, and 72°C for 1 min. Finally, we additionally incubated the tube at 72°C for 5 min.

We added 40 μL of Ampure XP beads to 50 μL of PCR solution. Purified sequence-library DNA was eluted into 20–30 μL of nuclease-free water. We checked the DNA concentration and DNA size of sequence library DNA using the Agilent High Sensitivity DNA Kit (Agilent) and QuantiFluor^®^ dsDNA System (Promega). We also prepared sequence library DNA with Tn5 transposase, in accordance with a modified version of a procedure used in a previous study [6]. Specifically, the modifications were as follows. We used 0.75 ng of amplified cDNA for the Nextera XT library preparation kit. We amplified sequence library DNA with the P5-gMac hybrid primer and the Nextera XT primer, which has the P7 sequence.

### Deep sequencing for Quartz-Seq2

We analyzed the sequence library DNA from Quartz-Seq2 using NextSeq500 (Illumina) and HiSeq2500 (Illumina). For the Read1 sequence, we used a custom sequence primer named Read1DropQuartz primer (HPLC-purified). In the case of the v3.1 RT primer, sequence specification was as follows (Read1: 22 cycles, Index1: 6 cycles, Read2: 64–118 cycles). In the case of the v3.2 RT primer, sequence specification was as follows (Read1: 23 cycles, Index1: 6 cycles, Read2: 63 cycles). When Read1 length was within 64 nt we mainly used the NextSeq 500/550 High Output v2 Kit (75 cycles), which can be used for sequencing up to 92 nt. For long sequences (over 64 nt), we analyzed sequence library DNA using the HiSeq Rapid SBS Kit v2. Sequence analysis with HiSeq2500 was supported by the Genomic Network Analysis Support facility (GeNAS) in RIKEN and Phyloinformatics Unit of RIKEN Center for Life Science Technologies (formerly GRAS) in RIKEN.

### Amplification-free single-cell qPCR analysis

Amplification-free single-cell qPCR was performed in accordance with our previous study with the following modifications [15]. We collected a single cell into 1 μL of lysis buffer (2.5 μM random hexamer, 1 mM dNTP mix, 0.3% NP40, 2 units/μL RNasin plus) of a 384-well PCR plate (Eppendorf) using SH800. The resulting 384-well PCR plate was cryopreserved at −80°C. We denatured total RNA in the 384-well PCR plate at 70°C for 1.5 min using a C1000/S1000 thermal cycler and hybridized the random primer to RNA at 0°C for 2 min. We added 1 μL of reverse-transcription solution (2x SSIV buffer, 10 mM DTT, 2 units/μL RNasin plus, 20 units/μL SuperScript IV) to the lysis buffer. Subsequently, we performed reverse transcription at 23°C for 10 min and 50°C for 10 min. The reverse transcription was stopped at 80°C for 10 min. The resulting solution was diluted with qPCR solution (10 mM Tris-HCl pH 8.0, 0.05% Tween-20). The obtained diluted solution was then used for qPCR detection with QuantiTect SYBR Green PCR Master Mix and the LightCycler480 system. For primer sets for each gene, see Additional file 3: Table S2.

### Drop-seq experiments and data analysis

Drop-seq was performed as reported previously [6] and in line with an online protocol (http://mccarrolllab.com/dropseq/), but with the following modifications: flow rates for oil and aqueous suspensions were 14,000 μl/h and 4,000 μl/h, respectively. The diameter of droplets was 95–100 μm. Microfluidic devices were fabricated by Fluidware Technologies (Japan). The lot number of barcoded beads was 051415 (ChemGenes). Data analysis for Drop-seq was performed as described online (http://mccarrolllab.com/dropseq/). The versions of the software and databases were as follows: STAR: v2.5.1b; mouse genome: GRCm38/mm10; genome annotation: gencode GRCm38.p4 vM9; and Drop-seq tools: v1.11.

### Quartz-Seq2 read alignment and generation of digital expression data

The structure of the sequence library and data processing was designed based on those of Drop-seq ([6] and online as referenced above). BCL files generated by Illumina NextSeq500 were converted to fastq files by bcl2fastq2 (v2.17.1.14) with demultiplexing pool barcodes. The --mask-short-adapter-reads parameter was set to 20. If needed, fastq reads were randomly downsampled by seqtk software (version: sgdp). We mainly trimmed read2 length to 62 nt for Quartz-Seq2 by using FASTX-Toolkit (version: 0.0.14). Fastq files for Read1 and Read2 were converted to a bam file by FastqToSam of Picard tools (version: 1.134). Extracting cell barcodes and UMI (also called as molecular barcode) and filtering out of reads with low barcode quality were performed using Drop-seq tools (version: 1.11). The resulting bam files were re-converted to fastq files by SamToFastq of Picard tools, and mapped to the mouse genome (GRCm38/mm10) using STAR (version: 2.5.1b). After sorting the resulting bam files using SortSam of Picard tools, the unaligned bam and aligned bam were merged by MergeBamAlignment of Picard tools. Then, gene names were assigned to each record using TagReadWithGeneExon of Drop-seq tools and gtf file (version: gencode GRCm38.p4 vM9). For the correction of errors of cell barcodes considering the Sequence–Levenshtein distance, a custom-made Python program (correct_bacode.py) was used, which enabled the correction of up to two nucleotide errors of substitution, insertion, or deletion. This program used Python2 (version: 2.7.11+), PypeR (version: 1.1.2), R (version: 3.2.3), Bioconductor (version: 3.2), and the Bioconductor package DNABarcodes (version: 1.0.0). Finally, the UMI for each gene for each cell was counted using DigitalExpression of Drop-seq tools, which generated the digital expression matrix. To generate a non-UMI-filtered matrix, a custom-made Python program that counts reads for each gene for each cell was used. To compare the quantitative performance between Quartz-Seq2 and reported data in Figure 4, we used Ensembl75 as a reference transcript. Moreover, we trimmed the read2 length to 45 nt for the comparison.

### Dimensionality reduction, clustering, and term analysis

Cells with low detected gene counts were removed for further analysis (Quartz-Seq2 on ES/PrE mixture: 4,000 genes; Quartz-Seq2 on SVF: 500 genes). Total counts of each cell were normalized to 10,000 or the mean of total UMI counts for cells. UMI counts had 1 added to them and were then log-transformed (natural logarithm), after which they were scaled to have mean and variance for each gene of 0 and 1, respectively. For PCA, UMI counts for all detected genes (for ES/PrE mixture) or genes with highly variable expression (for SVF) were used. For t-SNE, the top 10 to 40 principal components of matrices produced by PCA were used. For clustering, the DBSCAN algorithm was used. The values of the parameter epsilon were 0.69 for Quartz-Seq2 on the 4,500 mouse ES/PrE mixture, 5 for Quartz-Seq2 on the 384 mouse ES/PrE mixture, 3 for Drop-seq on the 500 mouse ES/PrE mixture, and 2.2 for SVF analysis. Significantly enriched GO terms with the top principal components were calculated using the GO-PCA package [45].

### Identification of differentially expressed genes for each cluster

Marker genes for each cluster were identified based on a generalized linear model. The difference in deviance between two models, in which the gene was or was not differentially expressed between two clusters, was calculated for the genes. To filter out noise, pseudocount 1 was added to the averaged gene expression in clusters and only genes with fold change of 2 or more between two clusters were further analyzed. P-values were calculated as 1 – cumulative density function of a chi-squared continuous random variable for difference in deviance. After corrections for multiple testing, genes with FDR of less than 0.05 were identified as differentially expressed genes for the cluster. For comparison between differentially expressed genes identified by scRNA-seq methods and those identified by bulk RNA-seq, this pseudocount was not added.

### Identification of variable genes in same cell-type

To identify genes for which the expression changes depending on the phase of the cell cycle, we used the intensity of Hoechst 33342 staining measured using a cell sorter. First, cells were discretized into 40 equal-sized buckets based on the rank of Hoechst 33342 staining intensity. Then, the CV of averaged UMI counts for a gene in each bin was calculated. After z-scaling of this, genes with high z-scores were identified. To identify genes whose expression fluctuates in a manner not relating to the cell cycle phase, we calculated the CV of UMI counts for a gene in each cell and the CV of averaged UMI counts for a gene in each bin. After z-scaling of such data, the genes for which the difference between two scaled CVs was large were identified as “variable genes the expression of which is less associated with the cell cycle phase”.

### Enrichment analysis on pathway and Gene Ontology (GO) terms

Pathways that were particularly enriched for the differentially expressed genes were calculated using the ReactomePA package of R Bioconductor (version: 1.14.4, [46]) and Metascape (http://metascape.org/ [47]). The cut-off parameter for the q value was 0.05. Terms that were enriched for genes with highly variable expression were calculated using DAVID, the Database for Annotation, Visualization and Integrated Discovery (version: 6.8, [48,49]). Ontology used for calculation was GOTERM_BP_FAT, GOTERM_MF_FAT, and GOTERM_CC_FAT. The terms with FDR < 0.05 were identified as enriched terms.

### Quantitative analysis for external RNA molecules

To calculate the ERCC capture efficiency, we determined the slope of regression between the input ERCC molecules and detected UMI counts for each ERCC. We tested our calculation using digital expression matrix (GSM1599501) of the inDrop single-cell sequence method as a control, and confirmed that our calculated ERCC capture efficiency (7.2%) approximately matched their reported ERCC capture efficiency (7.1%). To calculate the detection limit of ERCC molecules, the method of Svensson et al. was used [50]. We confirmed that our calculated results and those in the paper by Svensson et al. were approximately similar. For calculation of ERCC capture efficiency and detection limit for previous data, we used the concentration of ERCC spike-in RNA, which reported by original authors. For the paper by Ziegenhain et al., we confirmed the concentration by personal communication.

### Simulation-based power analysis

PowsimR was used to perform power analysis [51]. The parameters of the negative binomial distribution, namely, mean, dispersion, and dropout probability, were estimated by the function estimateParam with the following parameters: sigma = 1.96, distribution = “NB,” RNAseq = “singlecell,” and normalization = “scran.” To simulate differential expression between two groups using these parameters, the number of genes is represented by the number of detected genes (count > 0) in each method. Differentially expressed genes set up simulations with number of detected genes, 23.6% genes being DE, log fold change sample from a narrow gamma distribution (shape: 1.02, rate: 0.78). That parameter was derived from parameter estimates based on bulk RNA-seq data from mouse ES and PE cells (n = 3). The function simulateDE was used to run power analysis with the following parameters: DEmethod = MAST and normalization = scran. The results of power simulation were plotted using R with ggplot2 packages.

### Data resources

Raw and processed data files for Quartz-Seq2 and Drop-seq experiments are available under GEO: GSE99866 (https://www.ncbi.nlm.nih.gov/geo/query/acc.cgi?acc=GSE99866) for single-cell RNA-seq data and DRA002954 for bulk-RNA-seq data. A custom-made python program (correct_bacode.py) has been deposited under GitHub (DOI: 10.5281/zenodo.1118151, https://github.com/rikenbit/correct_barcode/).

## List of abbreviations

UMI: unique molecular identifier
PCR: polymerase chain reaction
PCC: Pearson’s correlation coefficient
SCC: Spearman’s rank correlation coefficient
CV: coefficient of variation
ES cell: embryonic stem cell
PrE cell: primitive endoderm cell
SVF: stromal vascular fraction
RT: reverse transcription
Dex: dexamethasone
PCA: principal component analysis
MSC: mesenchymal stem cell
TdT: terminal deoxynucleotidyl transferase
GO: Gene Ontology
FDR: false discovery rate
t-SNE: t-distributed stochastic neighbor embedding
nt: nucleotide
bp: base pair
SSC: side scatter
TPR: true positive rate
DEG: differentially expressed gene

## Declarations

### Ethics approval and consent to participate

Mouse experiments were carried out in accordance with the protocol approved by the National Institute of Advanced Industrial Science and Technology (AIST) Animal Care and Use Committee and the RIKEN Animal Experiment Committee.

### Consent for publication

Not applicable.

### Availability of data and material

The data have been deposited under GEO accession number GSE99866 and DRA accession number: DRA002954. A custom-made python program (correct_bacode.py) has been deposited under GitHub (DOI: 10.5281/zenodo.1118151, https://github.com/rikenbit/correct_barcode/).

### Authors’ contributions

YS, HD, and IN designed the study. YS designed and developed the Quartz-Seq2 reaction. YS and KT performed experiments of single-cell RNA-seq. HD, YS, and IN designed and performed data analysis. HT and AK designed and performed experiments on the mouse stromal vascular fraction. ME and KT assisted with the experiments. TH assisted with the development of the spin-down collection system. YS, HD, HT, AK, and IN wrote the manuscript. All authors read and approved the final manuscript.

## Acknowledgements

We greatly appreciate the Drop-seq team’s useful comments about our technical introduction to Drop-seq. We thank Mr. Akihiro Matsushima and Mr. Manabu Ishii for their assistance with the infrastructure for the data analysis. We are also grateful to Dr. Christoph Ziegenhain and Dr. Wolfgang Enard for giving us their quantitative data from single-cell RNA-seq methods. We also thank Dr. Rudolf Jaenisch and Dr. Yoichi Shinkai for giving us the J1 ES cell line. Advice and comments from Dr. Tempei Sato were also of great help in the culture of J1 ES cells. We thank Dr. Yutaka Suzuki for his assistance with the bulk RNA-seq. We are also grateful to Dr. Mika Yoshimura and Dr. Haruka Ozaki (RIKEN) for their assistance with the bulk RNA-seq analysis. Moreover, we thank the staff at the Phyloinformatics Unit of RIKEN Center for Life Science Technologies (formerly GRAS) for their assistance with the HiSeq2500 sequencer. Our thanks also go to Sony’s cell sorter team for specially customizing the 384-well sorting mode of the SH800. Finally, we thank Edanz Group (www.edanzediting.com/ac) for editing a draft of this manuscript.

